# Genetic Background Dictates Aortic Fibrosis in Hypertensive Mice

**DOI:** 10.1101/727800

**Authors:** Bart Spronck, Marcos Latorre, Sameet Mehta, Alexander W. Caulk, Abhay B. Ramachandra, Sae-Il Murtada, Alexia Rojas, Chang-Shun He, Bo Jiang, Mo Wang, Matthew R. Bersi, George Tellides, Jay D. Humphrey

**Author notes:** Correspondence: B. Spronck, Ph.D., Department of Biomedical Engineering, Yale University, New Haven, CT 06520, USA, or, J.D. Humphrey, Ph.D., Department of Biomedical Engineering, Yale University, New Haven, CT 06520, USA.

## Abstract

Many genetic mutations affect aortic structure and function in mice, but little is known about the influence of background strain. We phenotyped aortas from C57BL/6J and 129SvEv mice before and after continuous infusion of angiotensin II (AngII) for two weeks, which elevated blood pressure similarly in both strains (1.34-fold vs. 1.32-fold, systolic). Excised thoracic aortas were characterized functionally using isobaric vasoactive and cyclic passive stiffness tests whereas immunohistological studies quantified altered medial and adventitial composition as well as the infiltration of pan-inflammatory CD45^+^ cells. Baseline aortic geometry, composition, and biomechanical properties were similar across strains, consistent with mechanical homeostasis. Yet, aortic remodeling in response to AngII-induced hypertension differed dramatically between strains, with gross maladaptive remodeling in C57BL/6J but not in 129SvEv mice. CD45^+^ cell density was markedly higher in C57BL/6J than 129SvEv aortas while vasoconstrictive responses to AngII were greater in 129SvEv than C57BL/6J both before and after hypertension; importantly, smooth muscle mediated vasoconstriction reduces pressure-induced wall stress. Bulk RNA sequencing, layer-specific biomechanical modeling, and growth and remodeling simulations support the emergent hypothesis that mechanical stress-mediated immune processes promote maladaptive remodeling while smooth muscle contractile processes reduce wall stress and thereby protect against fibrosis. Differentially expressed mechano-sensitive genes thus play key roles in the distinct hypertensive aortic remodeling in C57BL/6J and 129SvEv mice and must be considered when comparing studies in different background strains, particularly mixed strains that are often used to generate mice with targeted mutations.

**Graphical Abstract:** 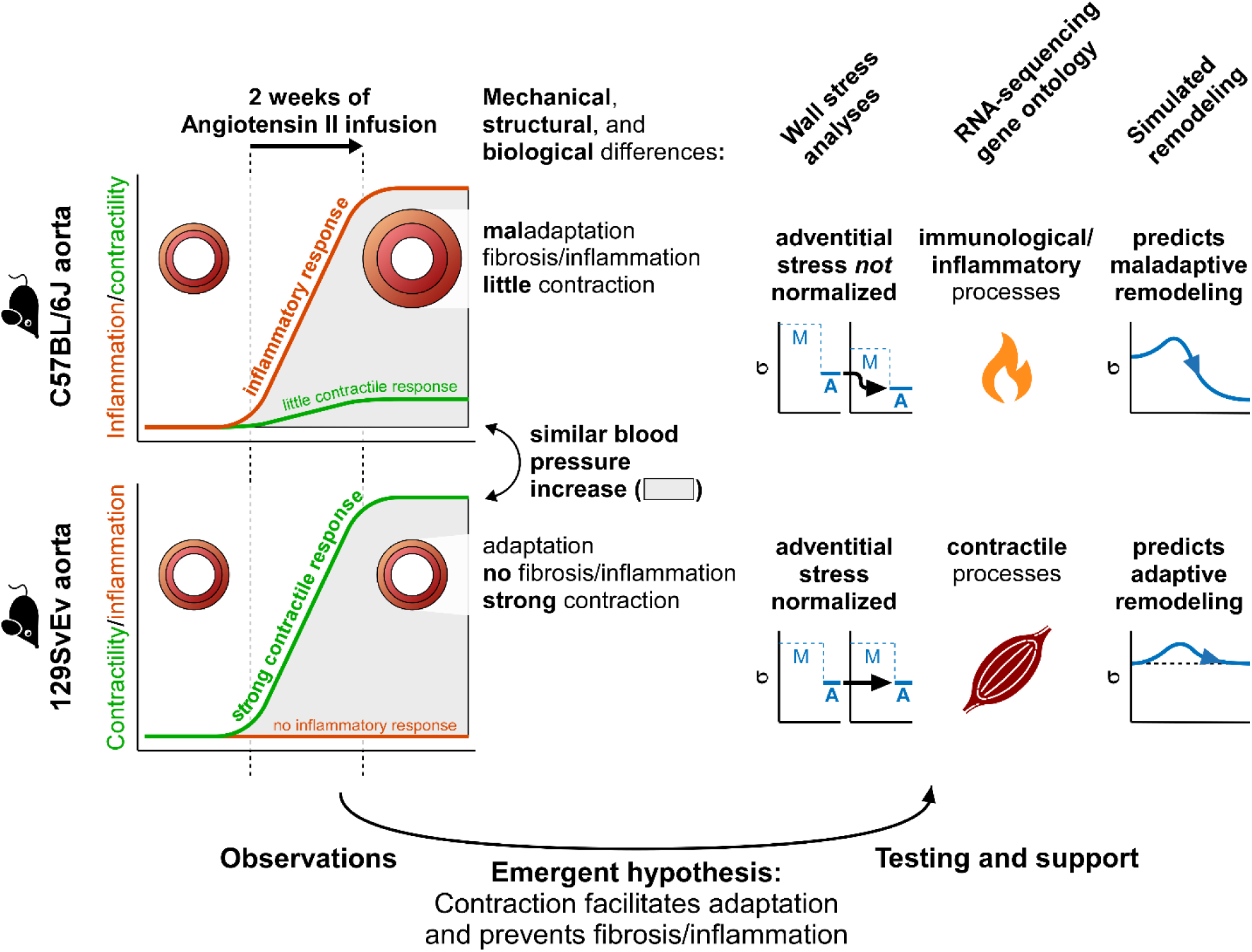

## Introduction

Mice have emerged as the most common animal model in vascular research for many reasons, including availability of antibodies for biological assays, ease of genetic manipulation, and short gestational period. Countless models are available, many of which exist on different genetic backgrounds. Pure backgrounds can reduce biological variability and enable statistical comparisons using smaller numbers of animals consistent with “reduction” in the three R’s of animal research. Conversely, mixed backgrounds can exhibit more variability as in natural populations, which will become increasingly relevant as we embrace precision medicine. The importance of genetic background is underscored further by the current lack of inclusion of diverse ancestral populations in human genomics research.^1^ Hence, the availability of diverse mouse models on different backgrounds promises to provide important information, though a challenge to analysis and interpretation. There is a pressing need for comparative studies.

Vascular phenotype is often described in clinical terms, including aneurysmal, atherosclerotic, fibrotic, stenotic, or tortuous. There is also a need to phenotype vessels biomechanically,^2^ however, given that mechanics governs many local cellular responses via the mechanobiology and many global circulatory responses via the physiology.^3-5^ Metrics for biomechanically phenotyping the aorta include multiaxial wall stress and material stiffness, the capacity to store energy elastically, and the ability to vasoconstrict or vasodilate under in vivo relevant biaxial loading conditions. We have quantified biomechanical differences in these functional metrics across both sexes and multiple regions of the central arterial system.^2,6,7^ Herein, however, we focus on male mice, for which more data are available for comparison, and one region, the descending thoracic aorta (DTA), which has a propensity for adverse remodeling in hypertension. Many reports similarly reveal biomechanical differences as a function of genetic mutations.^8-13^ Herein, however, we focus on differences due to background strain, which have not been studied in detail for the aorta. Specifically, we report marked differences in gene expression and biomechanical phenotype between two common strains of wild-type mice, C57BL/6J and 129SvEv, in response to hypertension induced by angiotensin II (AngII). Direct comparisons with prior data from other mice, including those on a C57BL/6;129SvEv background, provide further insight into the roles of genetic background in driving aortic fibrosis in hypertension.

## Methods

### Animals

Adult mice were obtained from Jackson Laboratory and Taconic Biosciences. Blood pressure was measured using a standard tail-cuff before and after infusion of AngII at 1000 ng/kg/min for two weeks using subcutaneous osmotic mini-pumps. Following euthanasia, four groups of DTAs were harvested for study ex vivo: normotensive (control) and hypertensive (AngII-infused) C57BL/6J and 129SvEv. All animal protocols were approved by the Yale University IACUC and conformed with NIH guidelines. Additional methods are in Supplemental Digital Content 1, including those describing the biaxial biomechanical testing, quantitative histology and immunohistochemistry, and bulk RNA sequencing as well as novel computational modeling of biomechanical metrics useful for assessing local mechano-adaptations or the lack thereof.

### Statistics

Data are presented as means±standard errors and analyzed using one- or two-way analysis of variance (ANOVA) as appropriate, with post-hoc Bonferroni tests; *p* <0.05 was considered significant. Differential gene expression was determined by bulk RNA-sequencing and considered for multiple comparisons using the Benjamini-Hochberg method.

## Results

### Baseline Mechanical Function is Similar Across Background Strains

Of the many possible biomechanical metrics, circumferential material stiffness and elastically stored energy are particularly important indicators of mechanobiological regulation of extracellular matrix and mechanical functionality, respectively.^2,9^ When evaluated at normal systolic pressures, values of mean circumferential material stiffness (1.90±0.09 MPa for DTAs from C57BL/6J mice and 1.85±0.05 MPa for those from 129SvEv mice) and mean elastic energy storage (79±4.1 kPa and 74±2.6 kPa, respectively) were not different statistically. Figure S1a-c (in Supplemental Digital Content 1) shows specimen-to-specimen comparisons of these key passive metrics as well as additional geometric and biomechanical metrics for both normotensive groups. Note the general consistency in all metrics across these two inbred strains, which is revealed further in Supplemental Table S1. Individual sample (raw) biomechanical data are provided in Supplemental Digital Content 2.

### Baseline Contractile Properties Differ by Strain

The ability of an artery to vasoconstrict against a constant luminal pressure at its in vivo axial stretch is an excellent indicator of smooth muscle functionality.^14^ Figure 1A shows normalized vessel-level vasoactive responses by control DTAs during isobaric (fixed pressure of 90 mmHg) and axially isometric (fixed specimen-specific in vivo axial stretch) studies ex vivo. The percent reduction in outer diameter and associated reductions in mean circumferential wall stress were similar for C57BL/6J and 129SvEv aortas (see, too, Figure S2) for both membrane depolarization (100 mM KCl) and *α*_1_-adrenergic stimulation (1 μM phenylephrine). Yet, there was a statistically greater vasoconstrictive response to an ex vivo bolus of 1 μM AngII by DTAs from control 129SvEv relative to C57BL/6J mice (−11% vs. -4% change in diameter, *p*=0.020; -24% vs. -8% change in circumferential stress, *p*=0.023; Table S2). There was no difference in maximum endothelial function, however, as revealed by changes in diameter resulting from ex vivo exposure to 10 μM acetylcholine, which promotes nitric oxide release by endothelial cells, or 1 mM L-NAME, which inhibits endothelial nitric oxide synthase (Figure 1A; Figure S2 and Table S2).

**Figure 1.**
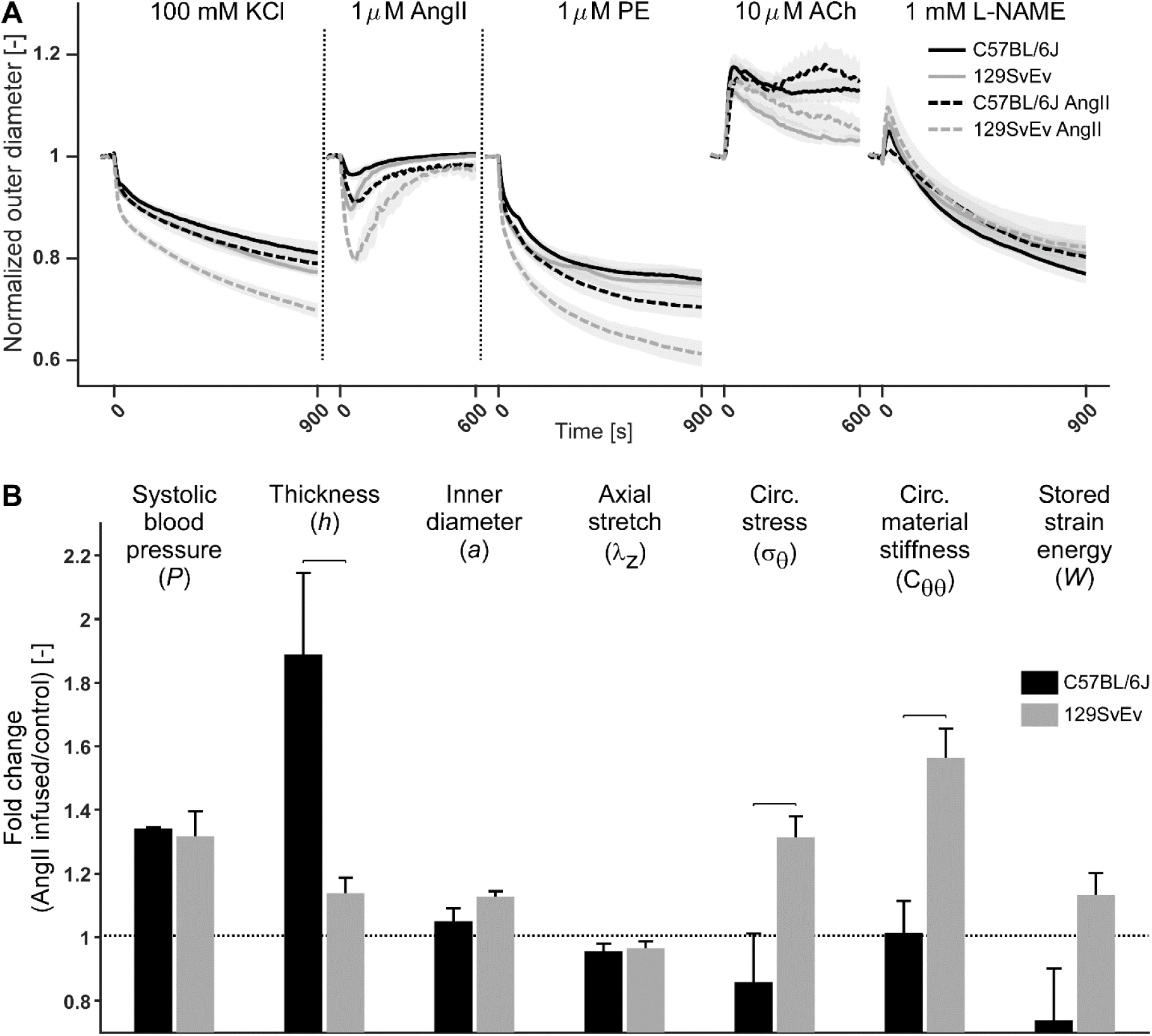
**A**: Thoracic aortas from 129SvEv mice show greater vasoconstriction to potassium chloride (KCl), angiotensin II (AngII), and phenylephrine (PE) than those from C57BL/6J mice and this difference is increased after AngII-induced hypertension; in contrast, vasodilatation is affected modestly by hypertension in both strains. Plots show normalized outer diameter as a function of time during ex vivo vasoactive testing for all four groups. Lines indicate mean values; shaded areas indicate the standard error. Vertical dotted lines indicate 600-second washout steps. Actual outer diameter responses are in Figure S3; values and statistics are in Table S2. ACh, acetylcholine; L-NAME, Nω-nitro-L-arginine methyl ester. **B**: AngII-induced fold-changes in passive biomechanical metrics: systolic pressure (*P*), deformed wall thickness (*h*), and deformed inner radius (*a*), each normalized by original normotensive values *P*_0_, *h*_0_, and *a*_0_. Note the larger fold change in *h* than in *P* in the C57BL/6J aorta, indicating maladaptation; conversely fold changes are much closer in the 129SvEv aorta. Also shown are in vivo axial stretch (*λ*_*z*_) and computed values of the transmurally averaged circumferential Cauchy stress (*σ*_θ_), circumferential material stiffness (*C*_θθ_), and elastically stored energy (*W*), all at systole. Overbars denote *p* < 0.05 (two-sample t-test).

### Passive Biomechanical Responses Differ by Strain in Hypertension

Notable differences manifested in the passive mechanical behaviors of hypertensive DTAs (Figure 1B) despite similar elevations in blood pressure: 1.34-fold increase for C57BL/6J (*P*_*sys*_ =174 mmHg relative to a baseline of 130 mmHg) and 1.32-fold for 129SvEv (*P*_*sys*_ =175 mmHg relative to 133 mmHg) mice (see, too, Figure S1 and Table S1). Notwithstanding similar reductions in the in vivo axial stretch (4%) in hypertension in both strains, the DTAs dilated less on average in C57BL/6J (5%) than in 129SvEv (13%) mice but thickened much more dramatically in C57BL/6J (89%) than in 129SvEv (12%) mice. Given the similar cardiac output in normotension and AngII-induced hypertension^15^ (i.e., *ε* = *Q*/*Q*_h_∼1, where *Q* is blood flow and subscript h denotes homeostatic), and similar fold-increases in blood pressure in hypertension across strains (*γ* = *P*/*P*_h_∼1.33), one would expect^16^ luminal radius *a* to remain nearly the same (*a* → *ε*^1/3^*a*_h_ = *a*_h_) while wall thickness *h* would increase proportional to the fold-increase in pressure (*h* → *ε*^1/3^*γh*_h_ = *γh*_h_) in a perfect mechano-adaptation. Hence, the gross over-thickening of the DTA in hypertensive C57BL/6J mice (89%, not an optimal 34%) is maladaptive, which reduced passive values of wall stress and energy storage well below normal (Figure 1B, Table S1). In contrast, the modest thickening of the DTA in hypertensive 129SvEv mice (12%, not an optimal 32%) appeared sub-optimal based on elevated passive values of wall stress and associated increases in material stiffness and energy storage at the new systolic pressure (Figure 1B, Table S1).

Stress, stiffness, and energy storage also increase with acute elevations in blood pressure due solely to a “pressure-effect” on a nonlinear material. This feature was evaluated using mechanical properties for the normotensive DTAs (Table S1) but for a simulated acute increase in pressure to the hypertensive level at a normal axial stretch (Figure S1c, grey rows labeled “@ higher *P*”). The calculated increases in stress, stiffness, and energy storage were only slightly higher for the normal 129SvEv aortas than those for the hypertensive ones, consistent with a sub-optimal adaptation of passive metrics to hypertension. In contrast, calculations for the C57BL/6J aortas revealed marked differences between measured and calculated values at the elevated pressure, consistent with a gross maladaptation in response to AngII-induced hypertension.

### Vasoconstriction Differs Dramatically by Strain in Hypertension

Whereas the vaso-responsiveness of the DTAs to 100 mM KCl and 1 μM PE increased modestly in C57BL/6J and 129SvEv mice following AngII infusion for two weeks, Figure 1A reveals further that the aforementioned greater responsiveness of the DTA from 129SvEv mice to an ex vivo exposure to 1 μM AngII both persisted and heightened following chronic AngII infusion in vivo. That is, the vasoactive reduction in diameter (Figure S2) and mean wall stress (Table S2) upon ex vivo stimulation with 1 μM AngII was significantly greater in 129SvEv than in C57BL/6J aortas following AngII infusion (−22% vs. -10% change in diameter, *p*<0.001; -46% vs. -24% change in stress, *p*<0.001).

Considering the significant contractions observed, and their differences between groups, we re-computed the mechanical metrics in Figure S1 for contractile states ranging from passive to maximally contracted (Figure S3). These results revealed that all but one 129SvEv DTA was able to restore circumferential wall stress to a normal (pre-infusion, normotensive) level through AngII-appropriate vasoconstriction in hypertension; the same was true for only one C57BL/6J DTA, however. Accounting for in vivo contractility suggests, therefore, that hypertensive remodeling in the 129SvEv DTAs was nearly optimal, whereas that for the C57BL/6J was grossly maladaptive.

### Immunohistological Characteristics Differ by Strain in Hypertension

Quantitative histology confirmed the gross thickening of the DTA in the hypertensive C57BL/6J mice (Figure 2A,B and Table S3), and localized it equally to the media (primarily increases in smooth muscle and collagen) and adventitia (increases in collagen). Immuno-histochemical staining for CD45-positive (CD45^+^) cells revealed marked increases in these inflammatory cells in hypertensive C57BL/6J DTAs (*p*<0.001), primarily in the adventitia, but not in 129SvEv DTAs (Figure 2C,D). Part of the percent increase in the C57BL/6J aortas could be explained by their lower CD45^+^ content under baseline normotensive conditions, yet absolute CD45^+^ area increased significantly with hypertension in C57BL/6J mice (Table S4). When considering specimen-to-specimen differences, the hypertensive remodeling reflected by wall thickening correlated well with CD45^+^ content (*p*=0.034) and the strength of both AngII-induced and maximal vasoconstriction ex vivo (*p*=0.002 and *p*<0.001), with vasoconstriction the stronger predictor statistically (Table S5).

**Figure 2.**
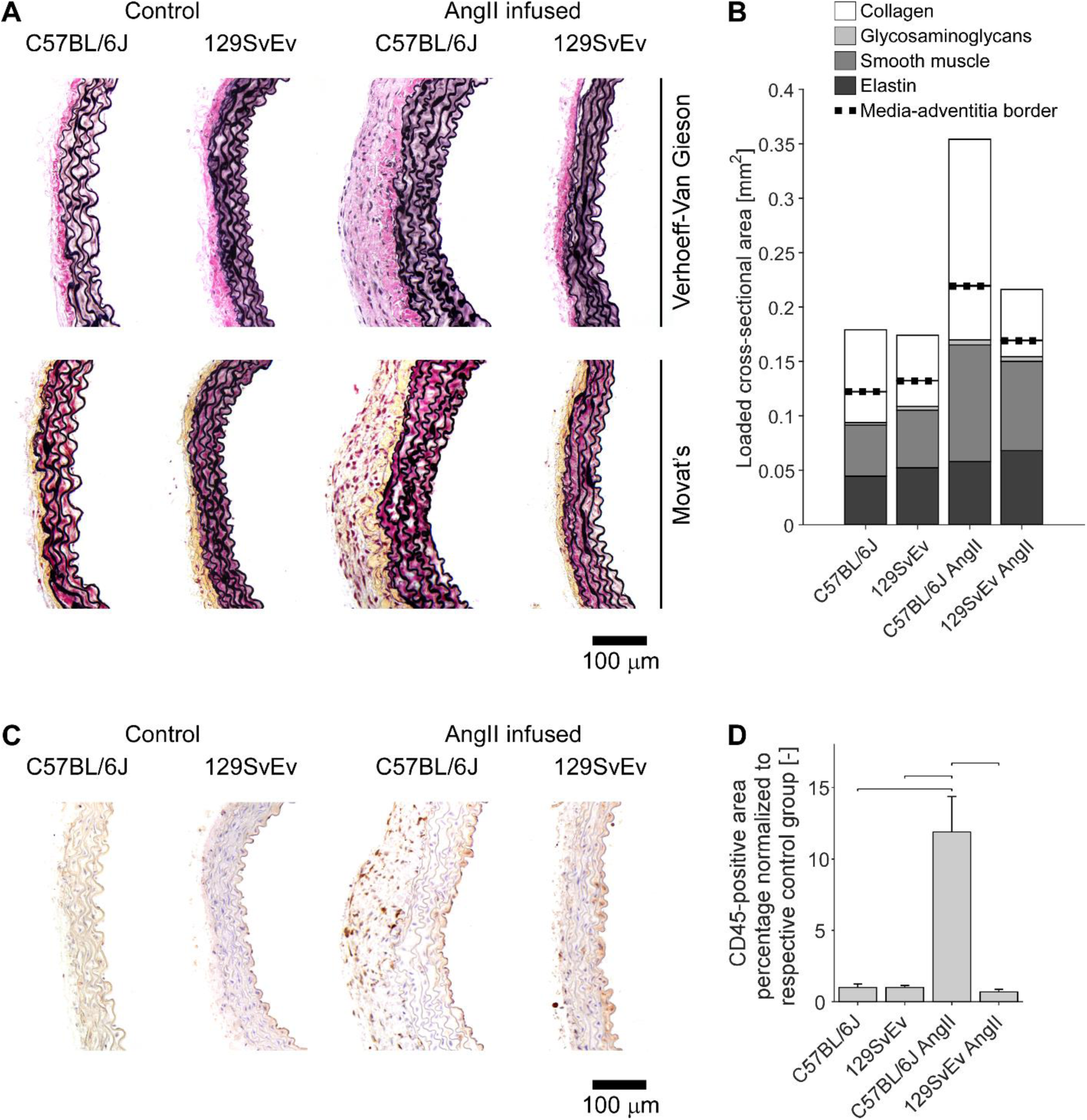
Chronic AngII-infusion causes marked medial and adventitial thickening and increased CD45 expression in thoracic aortas from C57BL/6J, but not 129SvEv, mice. **A**: Representative histological sections. **B**: Cross-sectional area of four major wall constituents for media (below dotted line) and adventitia (above dotted line). **C**: Representative CD45 immunohistochemistry. **D**: Normalized CD45^+^ cross-sectional area. Bars denote statistically significant differences (*p*<0.05, two-way ANOVA with post-hoc Bonferroni test). Quantification in Tables S3 and S4.

### Differentially Expressed Genes (DEGs) Differ by Background Strain in Hypertension

Consistent with the histo-mechanical assessments, bulk RNA sequencing (Figure S4) revealed that baseline gene expression was similar in the two wild-type groups; indeed, gene ontology (GO) analyses revealed only three GO categories that were mildly up-regulated in normal C57BL/6 relative to 129SvEv aortas (Figure S5a). Also consistent with the histo-mechanics, copious DEGs manifested when comparing hypertensive C57BL/6J and 129SvEv aortas (Figure S5b); for completeness, see differences in DEGs for the C57BL/6J DTAs before and after induced hypertension (Figure S5c) and similarly for the 129SvEv DTAs (Figure S5d). Many of the highly up- or down-regulated genes are of unknown biological significance (pseudogenes “*Gm…*”, Figures S5, S6), but Figure 3 shows maps of up- and down-regulated genes for the four study groups as a function of known phenotypic category (contractile, synthetic, immunity), with fibrillar collagen (*Col1a1, Col1a2, Col3a1, Col5a1, Col5a2, Col5a3*) and interleukin-6 (*Il6*) gene expression particularly high and smooth muscle contractile (*Acta2, Myh11, Cnn1*) gene expression low in the fibrotic AngII-infused DTAs from C57BL/6J mice. GO analyses further revealed numerous basic biological processes that differed between DTAs from hypertensive C57BL/6J and 129SvEv mice. Many of the top upregulated genes in the AngII-infused C57BL/6J aortas related to inflammatory and immune processes whereas many of the top upregulated genes in the AngII-infused 129SvEv aortas related to muscle contractility (Figure 4A,B; Figure S5). Full tables of all DEGs and GOs for all group comparisons are provided in Supplemental Digital Content 3. Finally, note that these strain-dependent differences in DEGs predicted well the strong ex vivo findings of roles of smooth muscle contraction protecting against and inflammation promoting aortic fibrosis (Figure 4C-E).

### Biomechanical Modeling Supports Experimental Findings

Notwithstanding consistent findings from bulk RNA sequencing and immuno-histo-mechanical data, it can be difficult to connect DEGs to overall functional effects. Hence, two computational biomechanical models were developed (Supplemental Digital Content 1) to explore further the emergent hypothesis that inflammatory cell infiltration drives a maladaptive, fibrotic aortic phenotype while smooth muscle cell contraction against exogenous AngII appears protective. First, we computed transmural distributions of wall stress with and without individually measured levels of contractile capacity. Under normal basal conditions, circumferential wall stress is borne largely by the media, which is expected teleologically because this layer contains the energy storing elastic laminae (Figure 2). This expectation was confirmed for DTAs from both strains under normotension for passive and multiple active states (Figure 5, and Video in Supplemental Digital Content 4). Importantly, simulated smooth muscle contractions against a bolus of 1 μM AngII, with an associated *γ* =1.33-fold elevation in pressure, enables medial and especially adventitial circumferential stress to return to within 10% of normal levels in the 129SvEv, but not C57BL/6J, DTAs.

**Figure 3.**
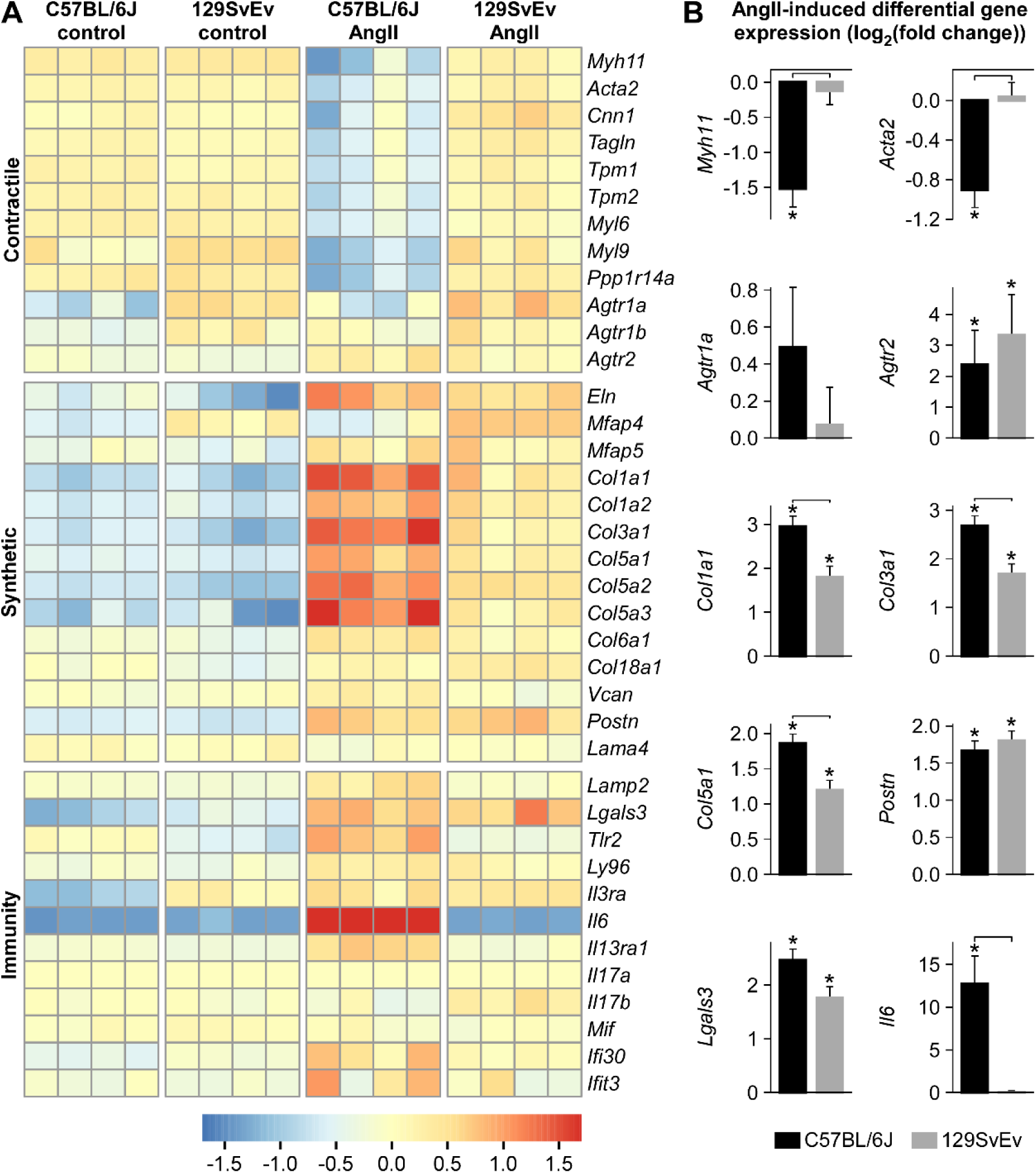
**A**: rlog-transformed gene expression for 16 individual samples across the four primary groups motivated by genes that drive smooth muscle cell phenotype^10^ — contractile, synthetic, or degradative. For each gene, mean rlog-transformed expression over all four groups was subtracted for normalization. Note, in particular, the downregulation of contractile markers and upregulation of matrix markers in the AngII-induced hypertensive C57BL/6J aortas. The gene encoding interleukin-6 (*Il6*) was significantly upregulated in this group as well; note that the color scale was capped at ±1.7 to avoid compression of all genes other than *Il6*, which showed absolute relative changes of 2.4 to 5.2. **B**: AngII-induced differential gene expression (after vs. before infusion). \**p*<0.05 for differential gene expression. Horizontal brackets indicate significant (*p*<0.05) differences between the differential gene expressions of the two strains (interaction effect). AngII, angiotensin II infused.

**Figure 4.**
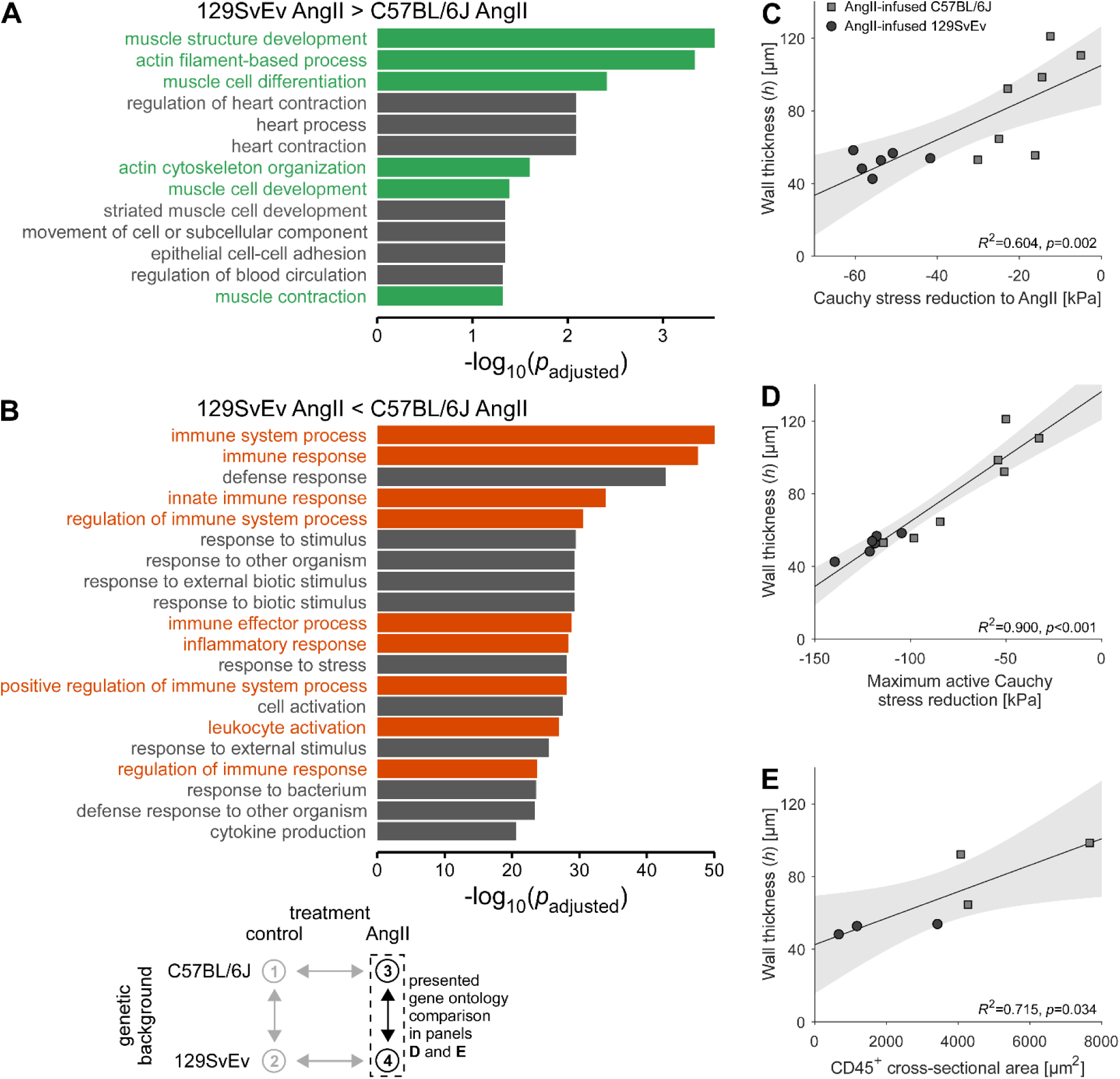
**A-B:** Bulk RNA sequencing-based gene ontology (GO) analyses revealed that biological processes associated with (**A**) muscle and its contraction were significantly upregulated in AngII-infused 129SvEv aortas relative to C57BL/6 aortas whereas those associated with (**B**) immune processes and inflammation were significantly upregulated in C57BL/6 aortas relative to 129SvEv aortas. **C-E:** Hypertension-induced changes in (**A**,**B**) wall thickness correlates negatively with the ability of the smooth muscle to contract in response to angiotensin II (AngII) and other stimulants but (**E)** correlates positively with CD45^+^ cell presence. Lines represent linear regressions, with 95%-confidence intervals shown with grey shading. Wall thicknesses were evaluated at a common pressure of 100 mmHg but individual axial stretches.

**Figure 5.**
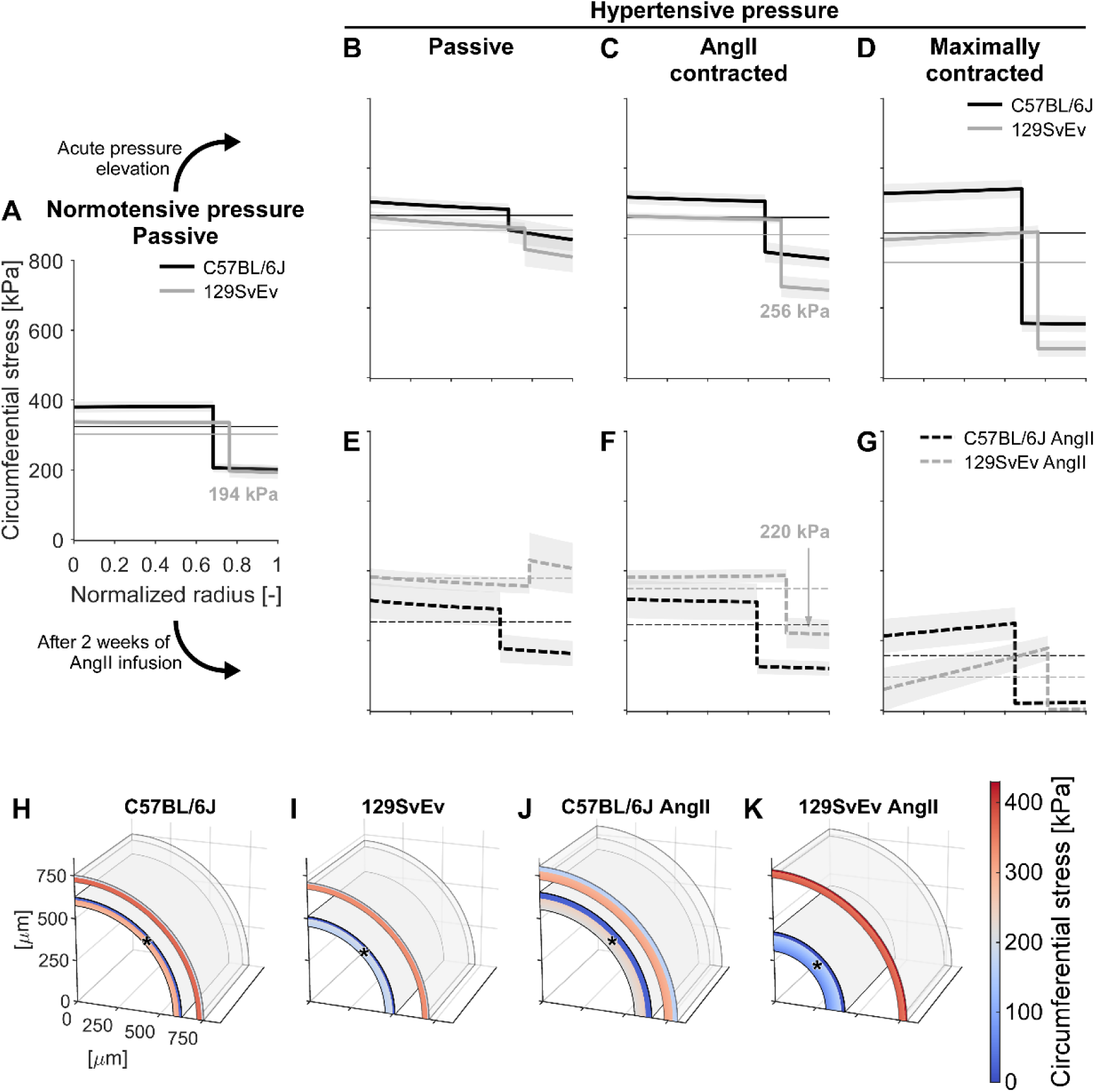
**A–G**: Calculated transmural distributions of circumferential Cauchy wall stress. Thin, horizontal lines represent the circumferential stress calculated using Laplace’s law. (**A**) The media carries most of the circumferential wall stress under normotensive conditions, as appropriate for storing elastic energy during mechanical loading that can be used during unloading to work on the distending fluid. (**B**) A simulated acute increase in blood pressure causes more load to be borne by the adventitia, (**C**) which can be reversed through vasoconstriction to angiotensin II (AngII). Panel **D** shows the vessel in a maximally contracted state. Note that even when maximally contracted, the C57BL/6J (C57) aorta — in contrast to the 129SvEv aorta — is unable to reduce its overall (Laplace) stress towards the normotensive value. (**E**) When only considering passive mechanical properties, AngII-induced hypertension and remodeling causes adventitial load bearing to surpass medial load bearing in 129SvEv but not in C57BL/6J (C57) aortas. (**F–G**) yet, the 129SvEv aorta is able to negate this *in vivo* through vasoconstriction. These simulations support the correlative finding of Figure 4 that increased smooth muscle contraction plays a key role in the appropriate remodeling of the aorta in the 129SvEv mice. Thin lines indicate mean values; shaded areas indicate the standard error. Panels **H–K** illustrate the large effect of maximum contraction (* denotes contracted state) on geometry and l stress; see the video in Supplemental Digital Content 4.

Second, we used a novel computational growth and remodeling (G&R) simulation (Supplemental Digital Content 1) to contrast dynamic processes of adaptation for DTAs from C57BL/6J and 129SvEv mice by predicting extracellular matrix production/removal in response to elevated blood pressure, both alone and in the presence of a wall stress-mediated inflammatory response. Figure 6 shows that, even when including AngII-appropriate levels of vasoconstriction, the predicted mean G&R of DTAs from the C57BL/6J mice resulted in excessive collagen deposition and hence wall thickening. In contrast, the predicted mean G&R for DTAs from the 129EvSv mice predicted near adaptation with AngII-appropriate levels of vasoconstriction. Additional simulations in Figure S7 with only basal smooth muscle contraction predict even higher collagen deposition and wall thickening for C57BL/6J DTAs and considerable wall thickening for 129EvSv DTAs. Both sets of simulations were thus consistent with experimental findings only when including a differential AngII-associated vasoconstriction that modulated inflammation via differences in the level of biaxial wall stress.

**Figure 6.**
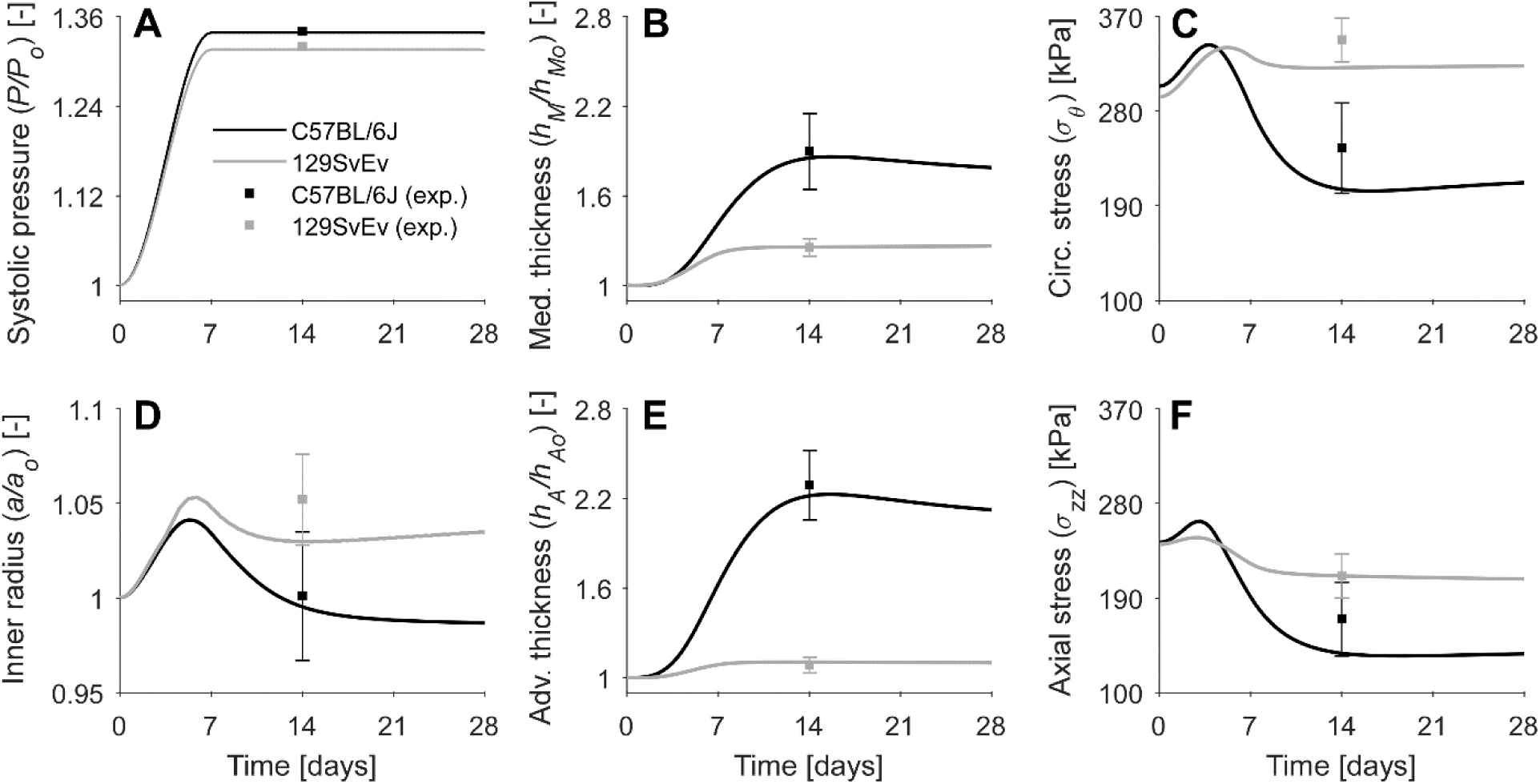
Growth & remodeling model predictions of responses following early, sustained 1.34- or 1.32-fold increases in systolic pressure (**A**, model input): (**D**) luminal radius, medial (med., **B**) and adventitial (adv., **E**) thickness (with quantities normalized to values at time 0 prior to AngII infusion) as well as mean circumferential (circ., **C**) and axial (**F**) stress for the C57BL/6J (black) and 129SvEv (grey) mice. Simulations are shown for a simultaneous increase in smooth muscle tone from day 0 (passive behavior) to respective AngII-appropriate levels under hypertensive conditions at day 7, then preserved. Stress-mediated inflammatory effects are computed internally by the model, resulting in a marked inflammatory response for the C57BL/6J but not the 129SvEv aorta (Figure S7, panel L). Shown, too, are means±standard errors of experimental values (squares with whiskers), noting that the model was informed by pre-hypertensive values, hence the simulations are predictions, not descriptions, of measured quantities. Note the dramatic increase in adventitial and medial thickness for the C57BL/6J aorta, and associated decrease in stress, by day 14, consistent with a delayed but exuberant production of inflammatory collagen primarily within the adventitia (Figure S7, panel G).

## Discussion

Many studies report differences across background strains in the (patho)biology of multiple organ systems in mice, including the vasculature (see Discussion in Supplemental Digital Content 1). We are not aware, however, of prior examination of the histo-mechanical phenotype of the aorta across background strains in induced hypertension, including passive and active biaxial responses that reflect structural and functional changes that result from specific DEGs. Differences herein between the hypertension-induced aortic thickening in C57BL/6J and 129SvEv mice were striking, and so too the associated consequences on wall stress, material stiffness, and energy storage. The DTA grossly maladapted in the C57BL/6J mice, consistent with prior studies,^17,18^ whereas the DTA appeared to under-adapt in the 129SvEv mice based on passive metrics but nearly adapted when considering smooth muscle tone. Importantly, these findings imply that prior observations of under-adaptation of passive aortic metrics in AngII-infused C57BL/6;129SvEv mice (Figure S8) likely arose from genetic modifiers associated with the 129SvEv strain, or more precisely from this specific sub-strain given the complex family tree of 129 mice.^19^ Indeed, a previously reported extreme maladaptation of the DTA in two of nine C57BL/6;129SvEv mice^7^ now appears to have been due to a greater penetrance of the C57BL/6 than the 129SvEv background in these two mice, noting that non-uniform segregation has been observed in intercrossed mice.^20^ Although archival tissue is not available to confirm this possibility, records indicate that the two maladaptive mice came from the same breeding pair while no other mice from that cohort descended from that particular litter. Hence, while we previously found that aging-related effects on biomechanical phenotype, especially luminal encroachment and reduction in material stiffness and elastic energy storage, are more severe than those of superimposed hypertension,^7^ it now appears that effects of genetic background can be even more important than effects of superimposed hemodynamic stress alone, at least when induced via chronic AngII infusion.

Clearly, the non-fibrotic aortic remodeling observed herein in response to chronic in vivo AngII infusion existed in vessels that had a persistently increased vasoresponsiveness to AngII ex vivo. Recall, therefore, that increased wall stress is a strong stimulant for wall remodeling,^3,5,16^ but vasoconstriction helps to reduce pressure-induced increases in wall stress since it decreases luminal radius *a* and increases wall thickness *h* isochorically (i.e., *σ*_*θ*_ = *Pa*(*P, C*)/*h*(*P, C*), where *P* is pressure, *C* is contractility). This important role of vasoconstriction is consistent with a recent finding in a larger cohort of C57BL/6 mice wherein a small sub-set of DTAs having modest AngII-induced hypertensive remodeling had a well-preserved vasoactive capability, at least to KCl.^21^ This observation is also consistent with our prior report of regional differences in AngII-induced hypertensive aortic remodeling in *Apoe*^*-/-*^ mice wherein maladaptive fibrotic remodeling manifested only in regions having a lower intrinsic vasoresponsiveness to AngII.^6^ That is, only the infrarenal aorta mechano-adapted in those mice (on a C57BL/6 background) and it alone exhibits strong constrictions in response to AngII in C57BL/6 mice (Figure S9). The magnitude of these contractions (12% reduction in diameter, 26% reduction in circumferential stress) is similar to the contractile response we observed herein for the DTA of normotensive 129SvEv mice (11% and 24% reductions, respectively). The increased contraction of the infrarenal aorta as compared to the DTA appears consistent with angiotensin II type 1 receptors existing at a higher density in the infrarenal than in the suprarenal, descending thoracic, and ascending aorta in C57BL/6 mice.^22,23^ We found greater AngII receptor expression in the DTAs of 129SvEv compared with C57BL/6J mice, though the greatest difference manifested in genes (e.g., *Myh11* and *Cnn1*) encoding smooth muscle-specific contractile proteins (Figure 3). Interestingly, aortic remodeling is modest-to-moderate in all regions in norepinephrine-induced hypertension,^24^ noting that these regions are vasoconstrictive to phenylephrine ex vivo in mice on a C57BL/6 background (Figure S9). There is clearly a need, therefore, to investigate further the role of local smooth muscle responsiveness to diverse exogenous stimuli in hypertension models, noting the often complementary roles of vasoactivity and matrix remodeling in arterial adaptations.^25,26^

Bulk RNA sequencing revealed many DEGs between the more fibrotic C57BL/6J and nearly adapted 129EvSv aortas. Importantly, both the GO analyses (Figure 4) and the biomechanical modeling studies (Figures 5,6) support the emergent hypothesis that local contraction against a exogenous vasoactive stimulus can reduce the pressure-induced intramural stress and thereby protect against inflammatory processes. This finding is consistent with a simpler analysis wherein comparisons of thoracic versus abdominal aortic remodeling in induced hypertension suggested that infiltration of inflammatory cells followed, rather than preceded, mechano-stimulated hypertensive remodeling and that inflammation drives fibrotic maladaptation.^6,17^

Many other factors likely play roles in regionally dependent vascular remodeling as well, including sex,^27^ regional differences in desmin/connexin expression^28^ and thus subtle differences in smooth muscle phenotype, local differences in pulsatile hemodynamic loading,^29^ and spatio-temporal differences in endogenous inflammatory burden.^6,17^ C57BL/6 mice have a high inflammatory propensity and are particularly susceptible to diet-induced atherosclerosis, obesity, and type 2 diabetes (e.g.,^30,31^). Hence, there is similarly a need to investigate further roles of inflammatory contributors to aortic remodeling in different hypertension models, noting that inflammation is a key driver of increased central artery stiffness in humans,^32,33^ which in turn increases pulse wave velocity and contributes to increases in central pulse pressure.^5^

Finally, there is also a pressing need to study in more detail the time-course of pressure elevation and associated aortic remodeling in both large and small vessels. Interestingly, large (elastic) arteries tend to remodel more in response to hypertension and aging than do medium-sized (muscular) arteries,^5^ which tend to have greater contractile responsiveness. Regarding time-course, we only studied DTAs after two weeks of AngII-induced hypertension in each model, thus we do not know whether DTA remodeling might have become maladaptive at longer times in the 129SvEv mice. We have shown previously, however, that AngII-induced hypertensive remodeling in C57BL/6 DTAs tends not to differ over two-to-four weeks of infusion^18,21^ and that hypertension induced in C57BL/6;129SvEv mice over longer periods (13 to 18 weeks) using a high salt diet and L-NAME did not cause adverse fibrotic remodeling in the DTAs.^7,34^ Hence, again, genetic modifiers intrinsic to the 129SvEv strain appear to have a dominant role in attenuating aortic remodeling in multiple models of induced hypertension in mice.

In conclusion, both transcriptional and functional data suggest a detrimental role of stress-mediated inflammation in promoting aortic fibrosis in (AngII-infused C57BL/6J mice) hypertension and conversely a protective role of smooth muscle contractility in reducing wall stress and thereby preventing fibrosis (in AngII-infused 129SvEv mice), both of which depend strongly on genetic background.

### Perspectives

- Baseline aortic geometry and properties are similar across pure (C57BL/6 and 129SvEv) and mixed (C57BL/6;129SvEv) background mice, suggesting common mechanical homeostatic targets;
- Aortic remodeling in response to chronic angiotensin II-induced hypertension differs dramatically between C57BL/6 and 129SvEv mice, suggesting a key role of genetic modifiers;
- Maladaptive aortic remodeling in hypertensive C57BL/6 mice associates with marked increases in CD45+ cells, suggesting a key role for stress-mediated inflammation; and
- Near adaptive aortic remodeling in angiotensin-induced hypertensive 129SvEv mice associates with greater aortic vasoresponsiveness to the exogenous AngII, suggesting an important role of contractility in reducing wall stress stimuli and preventing recruitment of inflammatory cells.

## Supporting information

Supplemental Digital Content 1

Supplemental Digital Content 2

Supplemental Digital Content 3

Supplemental Digital Content 4

## Author Contributions

Conceived Project (BS, MRB, GT, JDH), Performed Animal Experiments (CSH, BJ), Performed In Vitro Experiments and Modeling (BS, ML, AWC, ABR, SIM, MW), Analyzed Data (BS, SM, AR), Wrote Paper (BS, ML, JDH), Acquired Funding (BS, GT, JDH)

