## Supplemental Digital Content 1 for "Genetic Background Dictates Aortic Fibrosis in Hypertensive Mice"

[6]Vascular Biology and Therapeutics Program
Yale School of Medicine, New Haven, CT, USA

Correspondence:
B. Spronck, Ph.D.
Department of Biomedical Engineering
Yale University, New Haven, CT 06520, USA
+1-203-432-6678


or,

J.D. Humphrey, Ph.D.
Department of Biomedical Engineering
Yale University, New Haven, CT 06520, USA
+1-203-432-6428


**Supplemental Methods**

***Animals***. Adult male (14-20 weeks of age) C57BL/6J mice were obtained from The Jackson Laboratory (Bar Harbor, ME, USA) and 129SvEv (129S6/SvEvTac) mice were obtained from Taconic Biosciences (Rensselaer, NY, USA). Blood pressure was measured using a standard tail-cuff method (CODA, Kent Scientific, Torrington, CT, USA) while the mice were restrained within a transparent cylindrical tube without medication. Pressures were measured before and after infusion of angiotensin II (AngII; Sigma-Aldrich, St. Louis, MO, USA) at 1000 ng/kg/min for two weeks using an osmotic mini-pump (Alzet model #2004, DURECT Corporation, Cupertino, CA, USA) that was implanted subcutaneously on the flank while the mice were anesthetized with isoflurane (3% for induction, 1.5% for maintenance). Buprenorphine (0.1 mg/kg subcutaneously) was given pre- and post-operatively for analgesia. At the intended endpoint, the mice were euthanized with an intraperitoneal injection of pentobarbital sodium and phenytoin sodium (Beuthanasia-D; 150 mg/kg) and the descending thoracic aorta (DTA) was harvested and loose perivascular tissue removed gently. All biomechanical data collection and analysis was completed for $n$ = 6–8 mice for each of the four primary study groups: normotensive (NT) and hypertensive (HT) C57BL/6J and 129SvEv mice. Another $n$ = 5 mice per group were used for bulk RNA sequencing. All animal protocols were approved by the Yale University Institutional Animal Care and Use Committee (IACUC) and conformed to the current National Institutes of Health (NIH) guidelines.

To broaden the range of our comparisons, we also include historical passive biomechanical data from our laboratory for mixed background C57BL/6;129SvEv mice from two studies of genetic mutations, one for fibulin-5 ($Fbln5^{-/-}$) and one for smooth muscle myosin heavy chain ($Myh11^{R247C/R247C}$). Specifically, the experimental testing protocols and data analysis were identical for the following two mixed background strains: C57BL/6;129SvEv$_{Fbln5}$, which were obtained by breeding $Fbln5^{+/-}$ mice and selecting the $Fbln5^{+/+}$ mice, and C57BL/6;129SvEv$_{Myh11}$, which were obtained by breeding $Myh11^{+/R247C}$ mice and selecting the $Myh11^{+/+}$ mice. Further details on these mice and their phenotypes can be found elsewhere.[1,2]

***Biomechanical Testing.*** Excised vessels were mounted on custom drawn glass cannulae and secured at each end with 6-0 silk sutures. They were then placed within a custom computer-controlled biaxial device for biomechanical testing[3] and immersed in a heated (37°C) and oxygenated (95%/5% $O_2/CO_2$) Krebs-Ringer bicarbonate buffered solution containing 2.5 mM $CaCl_2$. The vessels were then subjected to a series of isobaric (luminal pressure of 90 mmHg) - axially isometric (fixed in vivo axial stretch) protocols wherein the vessels were contracted with 100 mM KCl, relaxed (KCl washed out), contracted with 1 µM AngII, relaxed (AngII washed out), contracted with 1 µM phenylephrine (PE), then dilated with 10 µM acetylcholine (ACh; without wash-out), an endothelial cell dependent stimulant of nitric oxide synthesis, and finally contracted with 1 mM $N_{\omega}$-Nitro-L-arginine methyl ester (L-NAME), a blocker of endogenous production of nitric oxide (a potent vasodilator).

Once these contraction-relaxation protocols were completed, the normal Krebs solution was washed out and replaced with a $Ca^{2+}$-free Krebs solution to ensure a sustained passive behavior while maintaining heating and oxygenation. Vessels were then preconditioned via four cycles of pressurization between 10 and 140 mmHg while held fixed at their individual in vivo axial stretch. Finally, specimens were exposed to three pressure-diameter (*P-d*) protocols, with luminal pressure cycled between 10 and 140 mmHg while axial stretch was maintained fixed at either the in vivo value or ±5% of this value, and to four axial force-length (*f-l*) tests, with force cycled between 0 and a force equal to the maximum value measured during the pressurization test at 5% above the in vivo axial stretch value, while the luminal pressure was maintained fixed at 10, 60, 100, or 140 mmHg. Pressure, axial force, outer diameter, and axial length were recorded on-line for all seven of these protocols and used for subsequent data analysis.

***Passive, Uni-layered Descriptors.*** We have found that five key biomechanical metrics define well the passive mechanical phenotype of the aorta: mean circumferential and axial wall stress, circumferential and axial material stiffness, and the elastically stored energy. Each of these metrics is calculated easily given best-fit values of the model parameters within a nonlinear stored energy function, which herein is assumed consistent with prior studies,[1,2,4-7] namely

$$W\left(\mathbf{C},\mathbf{M}^j\right)=\frac{c}{2}(I_{\mathbf{C}}-3)+\sum_{j=1}^{4}\frac{c_1^j}{4c_2^j}\left\{\exp\left[c_2^j\left(IV_{\mathbf{C}}^j-1\right)^2\right]-1\right\}, \tag{1}$$

where $c$ (dimension of kPa), $c_1^j$ (dimension of kPa), and $c_2^j$ (dimensionless, with $j=1,2,3,4$) are model parameters. $I_{\mathbf{C}}=\mathbf{C}:\mathbf{I}$ and $IV_{\mathbf{C}}^j=\mathbf{C}:\mathbf{M}^j\otimes\mathbf{M}^j$ are coordinate invariant measures of deformation, with $\mathbf{I}$ the identity tensor and $\mathbf{C}=\mathbf{F}^{\mathrm{T}}\mathbf{F}$ the right Cauchy-Green tensor, with $\mathbf{F}$ the deformation gradient tensor and the superscript $\mathrm{T}$ a transpose; $\det\mathbf{F}=1$ because of assumed incompressibility. The direction of the $j^{\mathrm{th}}$ family of fibers is given by the unit vector $\mathbf{M}^j=\left[0,\ \sin\alpha_0^j,\ \cos\alpha_0^j\right]$, with angle $\alpha_0^j$ computed with respect to the axial direction in a reference configuration. Based on prior microstructural observations, and the yet unknown effects of cross-links and physical entanglements amongst the multiple families of fibers, we included contributions of axial ($\alpha_0^1=0$), circumferential ($\alpha_0^2=\pi/2$), and two symmetric diagonal families of fibers ($\alpha_0^{3,4}=\pm\alpha_0$) to capture phenomenologically the complex biaxial material behavior; this relation has been validated independently.[8] The 3-D Cauchy stress tensor $\mathbf{t}$ is thus

$$\mathbf{t} = -p\mathbf{I} + 2\mathbf{F}\frac{\partial W}{\partial \mathbf{C}}\mathbf{F}^{\mathrm{T}}, \tag{2}$$

where $p$ is a Lagrange multiplier that enforces incompressibility; note that the transmurally averaged value of stress is denoted $\sigma$ by in the main text. Theoretical values of the applied loads were computed from components of Cauchy stress by solving global equilibrium equations in the radial and axial directions. Best-fit values of the eight model parameters were estimated via nonlinear regression (Levenberg-Marquardt) to minimize the sum-of-the-squared differences between experimentally-measured and theoretically-predicted values of luminal pressure and axial force, each normalized by average experimental measures.[1] Estimated parameters (constrained to be non-negative) were used to compute stress, material stiffness, and stored energy at any configuration. For example, components of the stiffness tensor ( $\mathcal{C}_{ijkl}$), linearized about a configuration defined by the systolic pressure and in vivo axial stretch, were computed as

$$\mathcal{C}_{ijkl} = \left. 2\delta_{ik}F^o_{lA}F^o_{jB}\frac{\partial W}{\partial C_{AB}} + 2\delta_{jk}F^o_{iA}F^o_{lB}\frac{\partial W}{\partial C_{AB}} + 4F^o_{iA}F^o_{jB}F^o_{kP}F^o_{lQ}\frac{\partial^2 W}{\partial C_{AB}\partial C_{PQ}}\right|_{\mathbf{C}^o}, \tag{3}$$

where $\delta_{ij}$ are components of $\mathbf{I}$, $\mathbf{F}^o$ is the deformation gradient tensor between the chosen reference configuration and a finitely deformed in vivo configuration, and $\mathbf{C}^o$ is the corresponding right Cauchy-Green tensor.

***Modeling Active Stress*** To study the effects of vasoconstriction on the aforementioned biomechanical metrics, we modeled active smooth muscle tone by assuming smooth muscle cells to develop a constant active second Piola-Kirchhoff stress, which corresponds to

$$W^{\mathrm{m,act}}(IV^{\mathrm{m}}_{\mathbf{C}}) = \frac{S_{\mathrm{act}}}{2}(IV^{\mathrm{m}}_{\mathbf{C}} - 1)\,, \tag{4}$$

where $S_{\mathrm{act}}$ denotes an active second Piola-Kirchhoff stress parameter for the medial layer, and $IV^{\mathrm{m}}_{\mathbf{C}} = \mathbf{C}:\mathbf{M}^{\mathrm{m}}\otimes\mathbf{M}^{\mathrm{m}}$ and $\mathbf{M}^{\mathrm{m}} = [0,\ \sin\alpha^{\mathrm{m}}_0,\ \cos\alpha^{\mathrm{m}}_0] = [0,1,0]$, assuming circumferential smooth muscle orientation ($\alpha^{\mathrm{m}}_0 = \pi/2$). In the presence of contraction, $W$ as used in Eqs. (2) and (3) is the sum of the passive (Eq. 1) and active (Eq. 4) contributions.

***Passive and Active Analysis of Bi-layered, Constituent-specific Transmural Wall Stress and Stiffness.*** In addition to the above analyses which do not mechanically distinguish the medial and adventitial layers, we performed layer-specific computations as introduced previously.[9] Briefly, we assumed constituent-specific strain energy functions and deposition stretches. Specifically, for elastin,

$$W^{\mathrm{e}}(\mathbf{C}^{\mathrm{e}}) = \frac{c^{\mathrm{e}}}{2}(I_{\mathbf{C}^{\mathrm{e}}} - 3)\,, \tag{5}$$

with $I_{\mathbf{C}^{\mathrm{e}}} = \mathbf{C}^{\mathrm{e}}\!:\mathbf{I}$, $\mathbf{C}^{\mathrm{e}} = \mathbf{F}^{\mathrm{e}^{\mathrm{T}}}\mathbf{F}^{\mathrm{e}}$, and $\mathbf{F}^{\mathrm{e}} = \mathbf{F}\mathbf{G}^{\mathrm{e}}_{\mathrm{h}}$, where $\mathbf{G}^{\mathrm{e}}_{\mathrm{h}}$ denotes the elastin deposition stretch tensor. For collagen family $j = [1,2,3,4]$,

$$W^{\mathrm{c}}\left(IV^{\mathrm{c},j}_{\mathbf{C}}\right) = \frac{c^{\mathrm{c}}_1}{4c^{\mathrm{c}}_2}\left\{\exp\left[c^{\mathrm{c}}_2\left(IV^{\mathrm{c},j}_{\mathbf{C}} - 1\right)^2\right] - 1\right\}, \tag{6}$$

with $IV^{\mathrm{c},j}_{\mathbf{C}} = \left(G^{\mathrm{c},j}_{\mathrm{h}}\right)^2\mathbf{C}\!:\mathbf{M}^{\mathrm{c},j}\otimes\mathbf{M}^{\mathrm{c},j}$ and $\mathbf{M}^{\mathrm{c},j} = \left[0,\ \sin\alpha^{\mathrm{c},j}_0,\ \cos\alpha^{\mathrm{c},j}_0\right]$. Similar to the uni-layered case, we let $\alpha^{\mathrm{c},1}_0 = 0$, $\alpha^{\mathrm{c},2}_0 = \pi/2$, and $\alpha^{\mathrm{c},[3,4]}_0 = \pm\alpha^{\mathrm{c}}_0$ a free parameter. Note that, in our bi-layered analysis, parameters $c^{\mathrm{c}}_1$ and $c^{\mathrm{c}}_2$ are assumed collagen family-independent. Deposition stretches $G^{\mathrm{c},j}_{\mathrm{h}}$, however, are collagen family-specific. For passive smooth muscle,

$$W^{\mathrm{m,pas}}(IV^{\mathrm{m}}_{\mathbf{C}}) = \frac{c^{\mathrm{m}}_1}{4c^{\mathrm{m}}_2}\{\exp[c^{\mathrm{m}}_2(IV^{\mathrm{m}}_{\mathbf{C}} - 1)^2] - 1\}\,, \tag{7}$$

with $IV^{\mathrm{m}}_{\mathbf{C}} = (G^{\mathrm{m}}_{\mathrm{h}})^2\mathbf{C}\!:\mathbf{M}^{\mathrm{m}}\otimes\mathbf{M}^{\mathrm{m}}$ and $\mathbf{M}^{\mathrm{m}} = [0,\ \sin\alpha^{\mathrm{m}}_0,\ \cos\alpha^{\mathrm{m}}_0] = [0,1,0]$, assuming circumferential smooth muscle orientation ($\alpha^{\mathrm{m}}_0 = \pi/2$). Active smooth muscle tone was modeled as in Eq. (4) and was assumed to only contribute to the medial layer. Hence, for the adventitial layer, $S_{\mathrm{act}} = 0$.

Total stored energy $W$ is now defined as

$$W = \varphi^{\mathrm{e}}W^{\mathrm{e}}(\mathbf{C}^{\mathrm{e}}) + \varphi^{\mathrm{c}}\sum_{j=1}^{4}\phi^{\mathrm{c},j}_{\mathrm{dir}}W^{\mathrm{c}}\left(IV^{\mathrm{c},j}_{\mathbf{C}}\right) + \varphi^{\mathrm{m}}[W^{\mathrm{m,pas}}(IV^{\mathrm{m}}_{\mathbf{C}}) + W^{\mathrm{m,act}}(IV^{\mathrm{m}}_{\mathbf{C}})], \tag{8}$$

with $\varphi^{\mathrm{e}}$, $\varphi^{\mathrm{c}}$, and $\varphi^{\mathrm{m}}$ the layer-specific constituent mass fractions, as estimated from quantitative histology (below), and $\varphi^{\mathrm{c},j}_{\mathrm{dir}}$ the (fitted) mass fractions of the different collagen fiber families, assuming $\sum_{j=1}^{4}\varphi^{\mathrm{c},j}_{\mathrm{dir}} = 1$ and $\varphi^{\mathrm{c},3}_{\mathrm{dir}} = \varphi^{\mathrm{c},4}_{\mathrm{dir}}$. Total Cauchy stress is then obtained using Eq. (2).

In describing our bi-layered mechanics, we assume equal constitutive parameter $c^{\mathrm{e}}$, $c^{\mathrm{c}}_1$, $c^{\mathrm{c}}_2$, $c^{\mathrm{m}}_1$, $c^{\mathrm{m}}_2$, and $\varphi^{\mathrm{c},j}_{\mathrm{dir}}$ values for the media and adventitia. Furthermore, to correctly capture non-tensile mechanics,[9] we assume a specific value for $c^{\mathrm{c}}_2$ and $c^{\mathrm{m}}_2$ under compression: $c^{\mathrm{m,c,comp}}_2$. Note that constituent mass fractions $\varphi^{\mathrm{e}}$, $\varphi^{\mathrm{c}}$, and $\varphi^{\mathrm{m}}$ *are* layer-specific.

Our parametrization approach deviates slightly from that described originally by Bellini and colleagues[9] in that (i) $c^{\mathrm{c}}_1$ and $c^{\mathrm{m}}_1$ were assumed to be valid under both compressive and tensile conditions (yielding a smooth stress response at the compressive-to-tensile transition), (ii) we estimated (fitted) instead of fixed the circumferential (2,2) component of $\mathbf{G}_\mathrm{h}^\mathrm{e}$, and (iii) we employed a single parameter estimation routine estimating all ten free passive ($S_\mathrm{act} = 0$) parameters simultaneously, which include $c^\mathrm{e}$, $c_1^\mathrm{c}$, $c_2^\mathrm{c}$, $c_1^\mathrm{m}$, $c_2^\mathrm{m}$, $c_2^\mathrm{m,c,comp}$, $\alpha_0^\mathrm{c}$, $(G_\mathrm{h}^\mathrm{e})_{2,2}$, $\varphi_\mathrm{dir}^\mathrm{c,1}$, and $\varphi_\mathrm{dir}^\mathrm{c,2}$ (note that $\varphi_\mathrm{dir}^\mathrm{c,3} = \varphi_\mathrm{dir}^\mathrm{c,4} = \left(1 - \phi_\mathrm{dir}^\mathrm{c,1} - \phi_\mathrm{dir}^\mathrm{c,2}\right)/2$). Deposition stretch values except $(G_\mathrm{h}^\mathrm{e})_{2,2}$ were assumed similar to those by Bellini and colleagues:[9] $(G_\mathrm{h}^\mathrm{e})_{3,3} = 1.6$, $(G_\mathrm{h}^\mathrm{e})_{1,1} = 1/\left((G_\mathrm{h}^\mathrm{e})_{2,2} \cdot (G_\mathrm{h}^\mathrm{e})_{3,3}\right)$; $G_\mathrm{h}^{\mathrm{c},j} = 1.1$ for all collagen families $j$; and $G_\mathrm{h}^\mathrm{m} = 1.1$.

After fitting the passive biomechanics and fixing the associated passive parameters, $S_\mathrm{act}$ was determined such that the modeled outer diameter matched either the outer diameter under passive or under contracted conditions.

***Local Mechano-Adaptations***. Copious studies suggest that arteries tend to mechano-adapt in response to modest changes in blood flow and blood pressure, that is, to maintain the flow-induced mean wall shear stress and pressure-induced mean circumferential wall stress near homeostatic targets.[10-12] If we let $\varepsilon$ denote the fold increase in blood flow ($\varepsilon = Q/Q_\mathrm{h}$) and $\gamma$ the fold increase in blood pressure ($\gamma = P/P_\mathrm{h}$), then it can be shown that a local mechano-adaptive response is given by $a \to \varepsilon^{1/3} a_\mathrm{h}$ and $h \to \varepsilon^{1/3}\gamma h_\mathrm{h}$, where $a$ is the current luminal radius and $h$ the current wall thickness, with subscript $\mathrm{h}$ denoting a homeostatic target value.[11,13] Note, too, that the mean circumferential stress can be evaluated as $\sigma_\theta = Pa(P,C)/h(P,C)$, which emphasizes that inner radius $a$ and wall thickness $h$ depend on both the distending pressure $P$ and the contractile state $C$ of the vessel, the latter of which is influenced by flow-induced changes in endothelial-derived vasoactive molecules, including nitric oxide and endothelin-1. Hence, the vasoactive state can influence dramatically the remodeling response.[10,14]

***Growth and Remodeling Model.*** We recently showed that a novel computational model of arterial growth and remodeling (G&R) that includes mechano- and immuno-stimulated matrix turnover can capture salient biomechanical features of the time-course of maladaptive remodeling of the thoracic aorta in C57BL/6 and *Apoe$^{-/-}$* (on a C57BL/6 background) mice infused with AngII.[15,16] Briefly, this constrained mixture model allows one to account for the evolution of mass fractions, mechanical properties, orientations, deposition stretches, and rates of turnover of multiple structurally significant constituents. Among the key equations is the mixture relation for biaxial wall stress

$$\boldsymbol{\sigma}_\Gamma^\alpha(s) = \frac{1}{\rho}\int_{-\infty}^{s} m_\Gamma^\alpha(\tau) q_\Gamma^\alpha(s,\tau) \frac{2}{\det \mathbf{F}_{\Gamma\,\mathrm{n}(\tau)}^\alpha(s)} \mathbf{F}_{\Gamma\,\mathrm{n}(\tau)}^\alpha(s) \frac{\partial \widehat{W}^\alpha\left(\mathbf{C}_{\Gamma\,\mathrm{n}(\tau)}^\alpha(s)\right)}{\partial \mathbf{C}_{\Gamma\,\mathrm{n}(\tau)}^\alpha(s)} \mathbf{F}_{\Gamma\,\mathrm{n}(\tau)}^{\alpha T}(s) d\tau \qquad (9)$$

with $\mathrm{n}(\tau)$ denoting potentially evolving constituent-specific natural (stress-free) configurations. Note the three primary constituent- ($\alpha = 1,2,\dots,n$) and layer- ($\Gamma = M, A$) specific constitutive functions in this constrained mixture formulation: the rate of mass production ($m^{\alpha}_{\Gamma}(\tau) > 0$, where $\tau$ is the time at which constituent $\alpha$ is produced), the removal of these constituents due both to their normal half-lives and accelerated losses in disease ($q^{\alpha}_{\Gamma}(s,\tau) \in [0,1]$, where $s$ is the current time of interest), and the passive multiaxial mechanical properties (captured by $\widehat{W}^{\alpha}(\mathbf{C}^{\alpha}_{\Gamma \mathrm{n}(\tau)}(s)) > 0$, which denotes the stored elastic energy in constituent $\alpha$ that depends on the deformation $\mathbf{C}^{\alpha}_{\Gamma \mathrm{n}(\tau)}(s) = \mathbf{F}^{\alpha T}_{\Gamma \mathrm{n}(\tau)}(s)\mathbf{F}^{\alpha}_{\Gamma \mathrm{n}(\tau)}(s)$ experienced by that constituent at current time $s$ relative to the configuration $\mathrm{n}(\tau)$) of the individual constituents. We let the stored energy function for elastin have a neo-Hookean form and for collagen and smooth muscle a Fung-type exponential form, which, together, constitute a layer-specific four-fiber family model. We let the rate of production of mass per unit reference volume be

$$m^{\alpha}_{\Gamma}(\tau) = m^{\alpha}_{\Gamma \mathrm{N}}(\tau)\Upsilon^{\alpha}_{\Gamma}(\tau) = m^{\alpha}_{\Gamma \mathrm{N}}(\tau)\left(1 + \mathrm{K}^{\alpha}_{\Gamma\sigma}\Delta\sigma(\tau) - \mathrm{K}^{\alpha}_{\Gamma\tau_w}\Delta\tau_w(\tau) + \mathrm{K}^{\alpha}_{\Gamma\varphi}\Delta\varrho_{\varphi}(\tau)\right) \tag{10}$$

where $m^{\alpha}_{\Gamma \mathrm{N}}$ are (potentially evolving) nominal values modulated by mechano- and immuno-biological stimulus functions $\Upsilon^{\alpha}_{\Gamma}$ that include deviations in mean pressure- and axial force-induced intramural stress $\Delta\sigma = (\sigma - \sigma_o)/\sigma_o$ and flow-induced wall shear stress $\Delta\tau_w = (\tau_w - \tau_{wo})/\tau_{wo}$ from homeostatic values ($\sigma_o$ and $\tau_{wo}$, each scalar metrics) as well as an inflammatory cell fraction $\Delta\varrho_{\varphi} \in [0,1]$ relative to its maximum possible density. Importantly, these three quantities are wall ( $\Delta\sigma$ and $\Delta\varrho_{\varphi}$) or luminal ( $\Delta\tau_w$) averages, with $\mathrm{K}^{\alpha}_{\Gamma\sigma}$, $\mathrm{K}^{\alpha}_{\Gamma\tau_w}$ and $\mathrm{K}^{\alpha}_{\Gamma\varphi}$ the constituent- and layer-specific gain-type parameters that modulate respective changes in cell/matrix production rate within each layer. Consistent with a first-order kinetic decay, we let

$$q^{\alpha}_{\Gamma}(s,\tau) = \exp\left(-\int_{\tau}^{s} k^{\alpha}_{\Gamma \mathrm{N}}(t)\,\mathrm{d}t\right) = \exp\left(-\int_{\tau}^{s} k^{\alpha}_{\Gamma o}\left(1 + \left(\Delta\sigma(t)\right)^2\right)\mathrm{d}t\right) \tag{11}$$

with the baseline rate parameter $k_{\Gamma o}$ modulated by the deviation in intramural stress $\Delta\sigma$. Finally, production and removal of constituent $\alpha$ determine its homogenized mass density per unit reference volume through

$$\rho^{\alpha}_{\Gamma \mathrm{R}}(s) = \int_{-\infty}^{s} m^{\alpha}_{\Gamma \mathrm{R}}(\tau) q^{\alpha}_{\Gamma}(s,\tau)\mathrm{d}\tau \tag{12}$$

where $m^{\alpha}_{\Gamma \mathrm{R}} = J_{\Gamma} m^{\alpha}_{\Gamma}$ are referential mass density production rates, with $J_{\Gamma} = \det \mathrm{F}_{\Gamma}$ the volume ratio computed from the layer-specific deformation gradient $\mathrm{F}_{\Gamma}$ from reference to current (in vivo) configurations for the mixture.

In previous studies,[15,16] the stress-driven evolution of inflammatory cell density in Eq. (10) was prescribed based on experimental findings (CD45+ staining). Here we describe it via a constitutive relation, similar to Eq. (12), that captures the main characteristics of the evolving infiltration of these cells, namely

$$\Delta\varrho_{\varphi}(s) = \int_0^s \mu_{\varphi}(\tau) q_{\varphi}(s,\tau) d\tau \tag{13}$$

with $\mu_{\varphi}$ a normalized infiltration term and $q_{\varphi}$ a survival function. In particular, we assume that there are no inflammatory cells at the onset of hypertension ($\Delta\varrho_{\varphi}(s=0)=0$). Removal is yet described by first order kinetics, with $q_{\varphi}(s) = \exp(-k_{\varphi}(s-\tau))$. For the rate of production / infiltration, we assume that the inflammatory response is triggered by a persistent level of high wall stresses, remaining active even if the stresses fall below the normal threshold during the subsequent (maladaptive) remodeling process. Hence, we model inflammatory cell infiltration as

$$\frac{d\mu_{\varphi}(s)}{ds} = \dot{\mu}_{\varphi}^{*} > 0 \quad \text{if} \quad \sigma \geq \sigma^{*} \quad \text{and} \quad \mu_{\varphi} < k_{\varphi} \tag{14}$$

where $d\mu_{\varphi}(s)/ds = 0$ otherwise, with $\dot{\mu}_{\varphi}^{*}$ a constant. Here, $\sigma^{*}$ represents a (scalar metric of) stress threshold level above which the infiltration of inflammatory cells into the tissue progressively increases; both parameters can be estimated from the known time course for the inflammatory response and the biaxial stresses (cf. Figures 2 and 5 in the main text). The second condition in Eq. (14) enforces a saturation value for $\mu_{\varphi}(=k_{\varphi})$ which will eventually lead to a saturation value for the normalized-to-maximum density $\Delta\varrho_{\varphi}(=1)$ via Eq. (13). Note that if $\sigma$ does not reach the inflammatory threshold $\sigma^{*}$ during the course of the hypertensive remodeling, then $d\mu_{\varphi}/ds = 0$, $\mu_{\varphi} = 0$, and $\Delta\varrho_{\varphi} = 0$, potentially leading to a purely mechano-adaptive remodeling in hypertension.[11] Finally, the value $\Delta\varrho_{\varphi}$, known from Eq. (13) at the current G&R time $s$, enters in the stimulus functions for smooth muscle and collagen production in Eq. (10) and, simultaneously modifies the inflammation-dependent passive properties in Eq. (9),[15,16] resulting in a coupled stress-driven immuno-mechanobiological response and an associated coupled G&R formulation.

***Quantitative Histology.*** Following mechanical testing, the aortas were fixed in their unloaded state in a 10% formalin solution for 24 h, stored in a 70% ethanol solution, embedded in paraffin, sectioned (5 µm thickness), mounted, and stained using three different stains: Movat's pentachrome (Mov), which stains elastin and nuclei black, collagen fibers yellowish, ground substance or glycosaminoglycans blue, and smooth muscle cells red; Verhoeff–Van Gieson (VVG), which stains elastin and nuclei black and collagen pink; and a cluster of differentiation 45 (CD45) antibody (ab10558, Abcam PLC, Cambridge, UK) at 1:100 with 3,3'-diaminobenzidine

(DAB) on a horseradish peroxidase (HRP) conjugated secondary antibody, which stains CD45-positive cells brownish-orange.

Images were acquired using an Olympus BX/51 microscope equipped with a DP70 digital camera (effective sensor resolution of $4080 \times 3072$ pixels, corresponding to a pixel size of $2.1\ \mu\mathrm{m}$ at a 2/3" ($8.8 \times 6.6$ mm) sensor size) and using a 20x objective (UPlanFl 20x, NA 0.50, optical resolution at $\lambda = 400$ nm of $\sim 0.49\ \mu\mathrm{m}$) and 0.5x tube lens, resulting in a total magnification of 10x and hence an image resolution of $0.21\ \mu\mathrm{m}$ (fulfilling the Nyquist criterion). Images were recorded using Olympus CellSens Dimension software. When arterial cross-sections exceeded the field-of-view, multiple images were acquired and stitched using Image Composite Editor software (Microsoft Research).

Three representative samples were selected for each of the four study groups, of which three sections were imaged for each of the three stains ($3 \times 4 \times 3 \times 3 = 108$ imaged sections). Custom MATLAB scripts were used to extract cross-sectional area fractions for elastin ($\varphi^{\mathrm{e}}$), collagen ($\varphi^{\mathrm{c}}$), smooth muscle ($\varphi^{\mathrm{m}}$) and ground substance/glycosaminoglycans ($\varphi^{\mathrm{g}}$). $\varphi^{\mathrm{e}}$ was averaged from VVG and Mov images; $\varphi^{\mathrm{m}}$ and $\varphi^{\mathrm{g}}$ were extracted from Mov images; $\varphi^{\mathrm{c}}$ was taken such that $\varphi^{\mathrm{e}} + \varphi^{\mathrm{c}} + \varphi^{\mathrm{m}} + \varphi^{\mathrm{g}} = 1$. CD45-positive area was similarly detected through hue-saturation-lightness thresholding.

***Bulk RNA Sequencing and Analysis***. Five additional mice per group (*n*=20 total) were used for RNA-Seq analysis. After euthanasia, crushed whole descending thoracic aorta (DTA) tissue was immersed in RLT lysis buffer (Qiagen N.V., Venlo, The Netherlands) and vigorously vortexed. Total RNA was isolated using a RNeasy Mini Kit and DNase Digestion Set (Qiagen) according to the manufacturer's protocol. For each group, the four samples with the highest RNA integrity number (RINe) were used for further analysis (*n*=16 total). Next-generation, whole-transcriptome sequencing was performed using a NovaSeq 6000 System (Illumina, Inc., San Diego, CA) at the Yale Center for Genome Analysis. RNA-Seq reads were aligned to a reference genome (GRCm38/mm10) with Gencode annotation[17] using HISAT2 for alignment[18], and StringTie for transcript abundance estimation.[18] Differentially expressed genes (DEGs) were obtained from raw gene counts using DESeq2 v1.26.0 in R v3.6.2.[19] Benjamini-Hochberg correction as implemented in DESeq2 was used to obtain p-values adjusted for multiple comparisons ($p_{\mathrm{adj}}$); genes with $p_{\mathrm{adj}} \leq 0.05$ were assumed to be differentially expressed. To reduce spurious large fold changes for genes with low read counts or high coefficients of variation, $\log_2$ fold change shrinkage was performed using the apeglm method.[20] Differentially expressed genes were used for gene ontology (GO) analysis for biological processes using goseq v1.38.0, run separately for up- and downregulated genes.[21] Benjamini-Hochberg correction was again used to adjust for multiple comparisons; categories with $p_{\mathrm{adj}} \leq 0.05$ were considered statistically significant.

***Statistics.*** Data were analyzed using one- or two-way analysis of variance (ANOVA) as appropriate, with post-hoc Bonferroni tests. Values are presented as means±standard errors. All mechanics and histology data processing and statistics, was performed in MATLAB R2019a (Mathworks, Natick, MA).

**Supplemental Discussion**

***Background Effects.*** There are many prior reports of differences in phenotype as a function of background strain. In the vasculature, capillary density increases more quickly in sv129 than in C57BL/6 mice following hind-limb ischemia;[22] mutations to the gene that encodes fibrillin-1 (*Fbn1*) and causes Marfan syndrome result in an earlier disease presentation in 129/Sv than in C57BL/6 mice;[23] consequences of elastin haploinsufficiency are greater in C57BL/6J;*Eln*$^{+/-}$x129X1/SvJ mice than in C57Bl/6J;*Eln*$^{+/-}$ mice;[24] and the atherosclerotic burden in the aortic arch of *Apoe*$^{-/-}$ mice is greater for 129SvEv than C57BL/6J mice.[25] We previously compared passive biaxial mechanical properties of the ascending aorta across seven different "wild-type" mice, including C57BL/6 and C57BL/6;129SvEv.[26] Few differences manifested except in non-induced mice that were modified for tamoxifen-induction of a particular genetic target.

Baseline blood pressures are higher in 129/SvJ than in C57BL/6 mice,[27] but the difference between 129SvEv and C57BL/6J mice (~2% systolic) was small herein. That 129/SvJ mice have two renin genes whereas C57BL/6 mice have one[27] could be important in AngII-infusion models, noting that plasma renin tends to be lower in 129/SvJ than in C57BL/6 mice and to decrease in AngII infusion (100-fold lower for 1000 ng/kg/min AngII infusion and 50-fold for 400 ng/kg/min), as one would expect of a system trying to reduce plasma AngII levels toward homeostatic values (~$3 \cdot 10^{-10}$ mol/L).[28] Whereas the luminal diameter of the DTA is similar in 129SvEv and C57BL/6J mice, the latter have a higher stroke volume and heart rate (and thus cardiac output). Yet, mean blood velocity is similar in the DTA and so too the mean wall shear stress in vivo.[25] We are the first to show quantitatively, however, that key passive geometric and mechanical metrics are similar for the DTA for these two strains as well as for two related mixed C57BL/6;129SvEv strains (Tables S1 and S6). We also found some similarities but also a key difference in the vasoconstrictive responses of the DTA between 129SvEv and C57BL/6J mice. Vasoconstriction was similar herein for phenylephrine and only slightly greater in the 129SvEv mice for KCl stimulation, but the ex vivo response to AngII was nearly three-fold greater in DTAs from normotensive 129SvEv than C57BL/6J mice and two-fold greater in hypertension (Table S2). These differences are critical given the widespread usage of AngII to induce hypertension and related pathologies in mice.[29-31] Similar findings have been reported for the mesenteric artery,[32] which is a muscular, not elastic, artery. Because arterial remodeling is influenced strongly by vasoactivity,[10,14] it is important to delineate smooth muscle responses across strains, which is seldom done. For example, Hussain et al.[33] did not detail potential differences in background in studies of the contractility of the mouse thoracic aorta from eNOS and nNOS null (on a hybrid C57BL/6;SV129 background) versus iNOS null (on a pure C57BL/6 background) mice.

***Consistent Comparisons with Prior Data.*** Passive baseline aortic properties for our two pure strains (C57BL/6J and 129SvEv – present data) and two mixed strains (C57BL/6;129SvEv$_{\mathit{Fbln5}}$ and C57BL/6;129SvEv$_{\mathit{Myh11}}$ – historical data)[2,7] are very similar (Figure S8). Note, therefore, that we have previously subjected C57BL/6J[5] and C57BL/6;129SvEv$_{\mathit{Fbln5}}$[7] mice to two weeks of AngII infusion at a lower rate (490 ng/kg/min). In response to 1.33- to 1.38-fold increases in blood pressure over two weeks, the DTA grossly over-thickened in the C57BL/6J mouse whereas it only thickened modestly in the mixed C57BL/6;129SvEv$_{\mathit{Fbln5}}$ mouse, the latter reminiscent of the response of the DTA to a higher rate of AngII infusion in 129SvEv mice (present study). Importantly, both the 490 and 1000 ng/kg/min rates of AngII infusion in the C57BL/6J mice yielded (on average) nearly equal fold increases in blood pressure, $\gamma \approx 1.34$ (telemetry measurement[5]) and 1.34 (tail-cuff measurement, present study), respectively. It thus appears that attenuated responses in the C57BL/6;129SvEv aorta to AngII-induced hypertension were dominated by the 129SvEv ancestry. For this reason, the elevated values of wall stress, material stiffness, and stored energy in the mixed background mice again likely reflected primarily a pressure-effect, namely pressure-dependent values for an aortic wall that is otherwise similar to that in the normotensive mouse. This also holds for the decreased values in distensibility, a metric of structural stiffness commonly used clinically. To assess this possible pressure dependence, we again computed values of stress, stiffness, and energy storage for the normotensive groups for a hypothetical acute increase in pressure to the corresponding chronic AngII-infused value (Table S7, "@ higher *P*" rows). In this mixed background-group, we again found that the predicted fold-changes were close to the measured fold-changes in the chronically infused group, suggesting modest passive wall remodeling in the C57BL/6;129SvEv mice. In contrast, predicted and measured fold-changes differed markedly in the lower rate of infusion C57BL/6J group, consistent with a maladaptive (over-) remodeling in these mice.

Finally, we compared the present findings with historic DTA data from our laboratory for AngII-infused *Apoe*$^{-/-}$ mice (Figure S8),[6] which are on a C57BL/6 background and often used in AngII infusion studies because of their propensity for atherosclerosis, aneurysm, or dissection. These mice were also subjected to two weeks of AngII infusion at the higher rate of infusion (1000 ng/kg/min), which can be compared directly to the data in Figure S1 for the pure C57BL/6J and 129SvEv mice. Interestingly, the *Apoe*$^{-/-}$ data reveal a greater increase in blood pressure over two weeks ($\gamma = 1.62$, tail cuff), yet a remarkable near perfect geometric mechano-adaptation (inner radius dilated only 3% whereas wall thickness increased 1.61-fold). The circumferential values of passive wall stress and material stiffness were nevertheless slightly higher, on average, than would be expected if fully mechano-adapted. Indeed, if one looks more carefully at the time-course of changes in the AngII-infused *Apoe*$^{-/-}$ DTA in the original report,[6] results at three and then four weeks of AngII infusion reveal a gross over-thickening of the wall, including a remarkable 3.85-fold increase for a $\gamma = 1.67$ fold increase in systolic pressure at four weeks. Hence, these later results are also generally consistent with the maladaptive results observed herein for a C57BL/6 background. Note that this delayed but exuberant over-thickening was ascribed to a similarly delayed infiltration of CD45$^+$ inflammatory cells, primarily in the adventitia where the fibrotic thickening occurred, also consistent with findings herein (Figures 5,6).

**Acknowledgments**

We are grateful for prior collaborations with Drs. Jacopo Ferruzzi, Hiromi Yanagisawa, Chiara Bellini, and Dianna M. Milewicz that yielded data for C57BL/6;129SvEv mice[1,2] and Dr. David G. Harrison for data for low-rate AngII infusion of C57BL/6 mice,[5] which were used herein as comparators.

**Supplemental Figures**

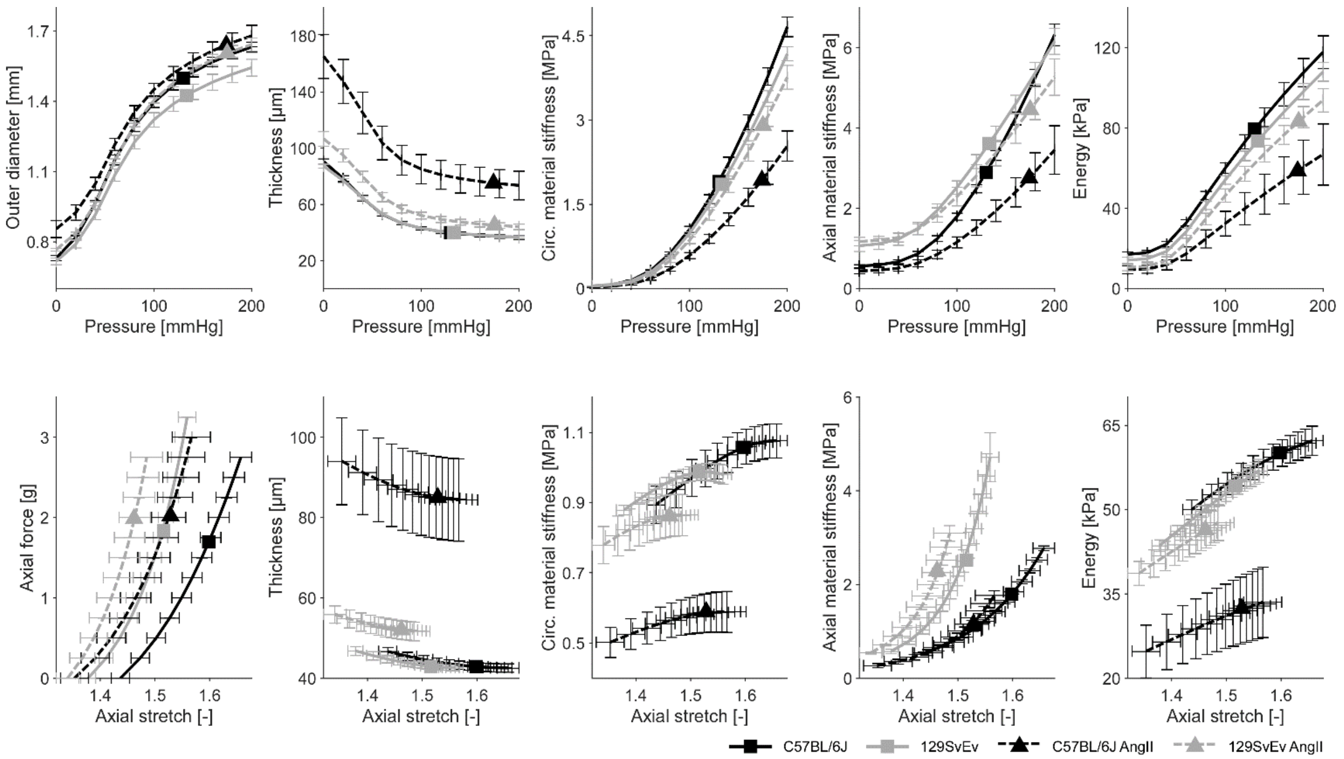

**Supplemental Figure S1a.** Fundamental passive biomechanical metrics such as wall stress, material stiffness, and elastic energy storage depend strongly on the biomechanical state, often defined conveniently by the distending pressure and axial stretch at which the metric is evaluated. Shown here are key metrics measured and/or calculated for the descending thoracic aorta based on data collected during the ex vivo biaxial tests for all four study groups: normotensive (control) and hypertensive (AngII-infused) C57BL/6J and 129SvEv. Metrics are plotted as a function of distending pressure for constant sample-specific in vivo values of axial stretch (top row) or as a function of axial stretch for a constant pressure of 100 mmHg (bottom row). Symbols indicate means, error bars indicate standard errors. Enlarged symbols represent values at systolic blood pressure (top row) and in vivo axial stretch (bottom row). See, also, Table S1 and additional data in Figures S1b and S1c.

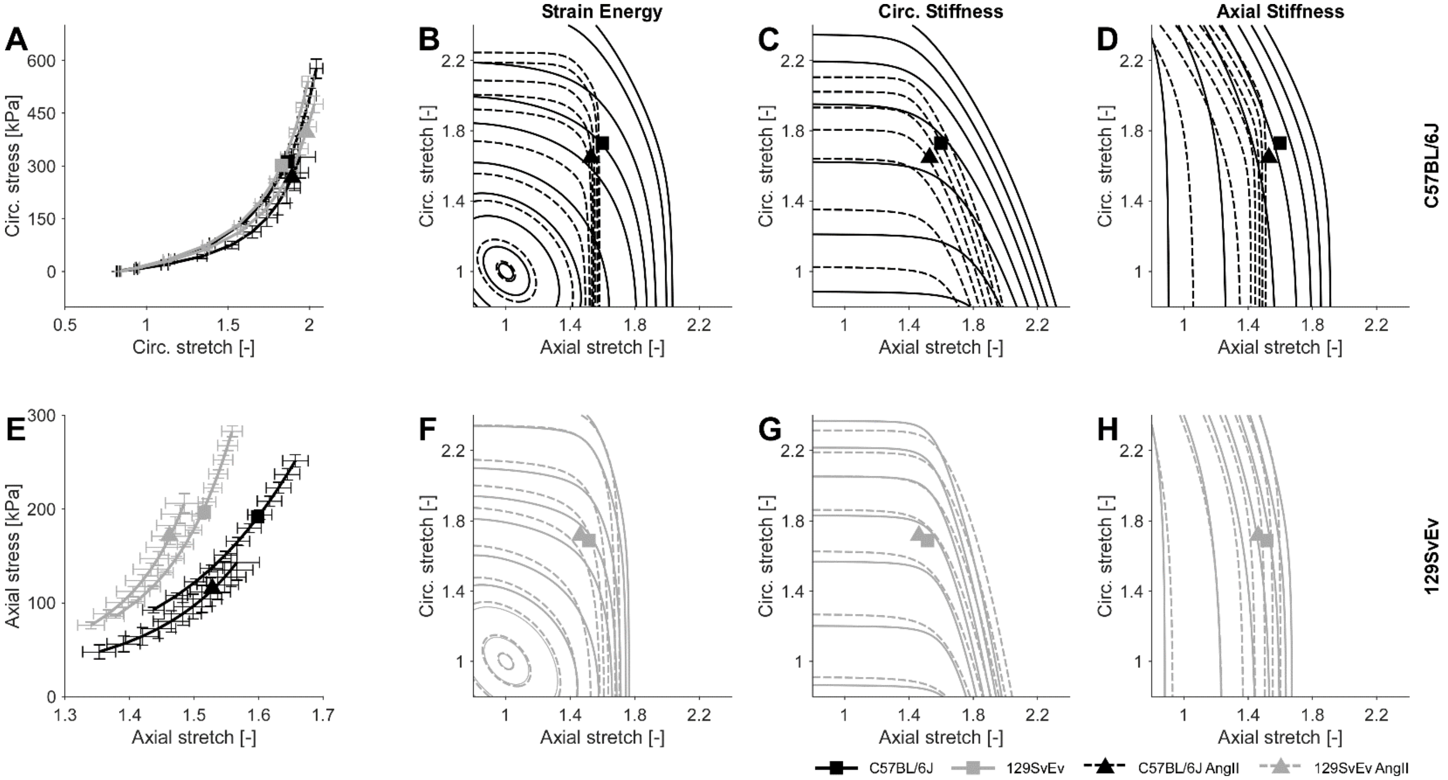

**Supplemental Figure S1b.** Panels A and E: passive circumferential and axial Cauchy stress-stretch curves for the descending thoracic aorta plotted at in vivo axial stretch (**A**) and a distending pressure of 100 mmHg (**E**) for all four study groups. Symbols indicate means, error bars indicate standard errors. Enlarged symbols represent values at systolic blood pressure (**A**) and at in vivo axial stretch (**E**). Panels **B**, **C**, and **D**: group-averaged contour plots of elastically stored energy, circumferential material stiffness, and axial material stiffness as a function of circumferential and axial stretch for normotensive (control – solid lines) and hypertensive (AngII-infused – dashed lines) C57BL/6J aortas. Enlarged symbols correspond to systolic blood pressure. Panels **F**, **G**, and **H** are the same as **B**, **C**, and **D** except for 129SvEv aortas. Notice the dramatic reduction in axial extension in C57BL/6J but not 129SvEv aortas. See, too, Figure S1c for sample-specific results. Raw data are provided in Supplemental Digital Content 2.

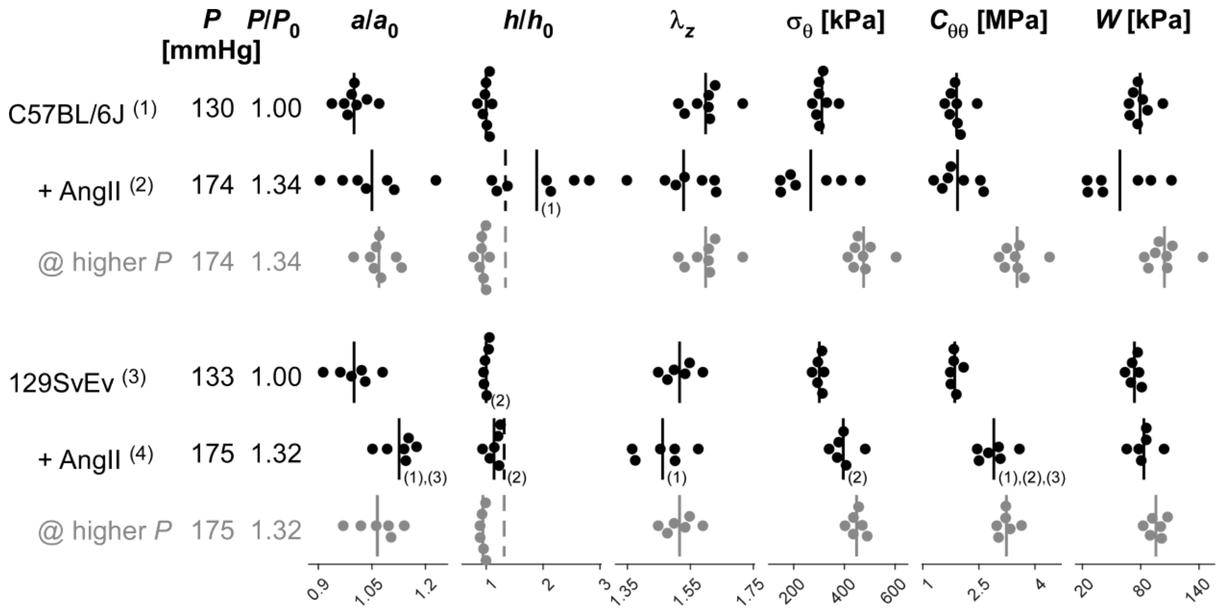

**Supplemental Figure S1c.** Specimen-specific values of passive biomechanical metrics for the descending thoracic aorta from all four study groups (1–4; black rows): normotensive (control) and hypertensive (+ AngII) C57BL/6J and 129SvEv mice. Shown are group-specific systolic pressure ($P$), deformed inner radius ($a$), and deformed wall thickness ($h$), each normalized by original normotensive values $P_0$, $a_0$, and $h_0$. Also shown are in vivo axial stretch ($\lambda_z$) and computed values of the transmurally averaged circumferential Cauchy stress ($\sigma_\theta$), circumferential material stiffness ($C_{\theta\theta}$), and elastically stored energy ($W$), all at systolic pressure. Vertical solid lines indicate arithmetic means; dashed lines in the $h/h_0$ column represent the ratio $\gamma = P/P_0$ to enable visual distinction of adaptive (lines overlap) versus under-adaptive (solid line to the left of the dashed line) or mal-adaptive (solid line to right of dashed line) passive remodeling. Grayed-out rows, labeled "@ higher *P*", show calculations for normotensive groups for an acute elevation of blood pressure that matches that for the AngII groups. Numbers in swarm plots indicate significant differences ($p < 0.05$) amongst the four groups, 1–4; no statistical tests were performed for the "@ higher *P*" rows. AngII, angiotensin II.

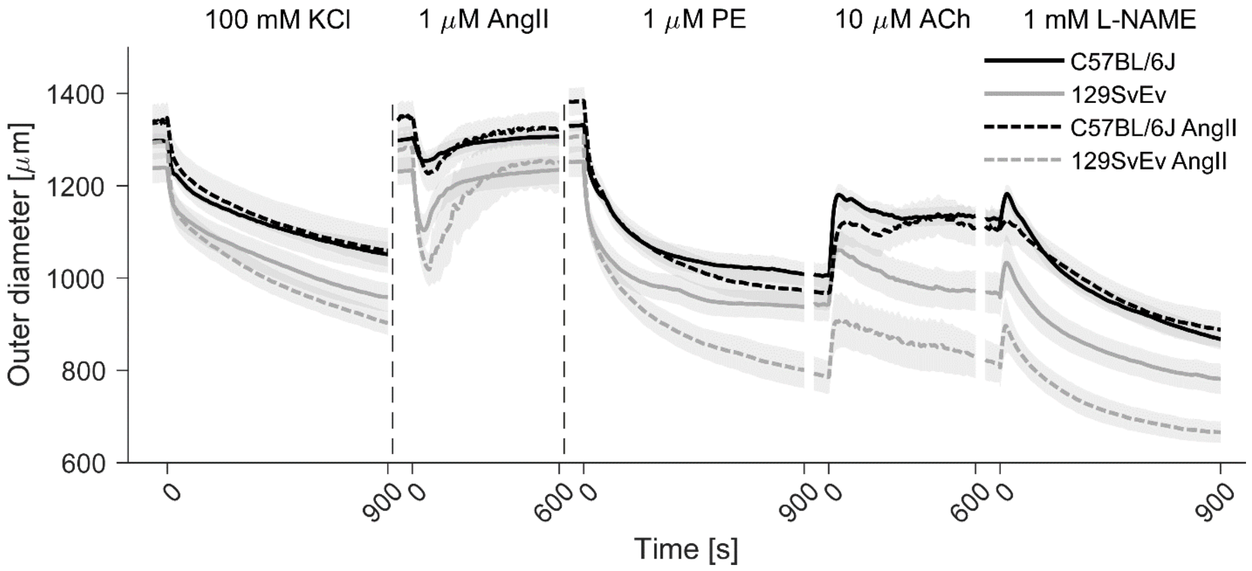

**Supplemental Figure S2.** Change in outer diameter (in microns) as a function of time during ex vivo vasoactive testing for all four study groups: C57BL/6J and 129SvEv without and with AngII-induced hypertension. The thick lines indicate mean values, shaded areas indicate standard errors. Vertical dashed lines indicate 600-second washout steps. Normalized outer diameter responses are shown in Figure 1 in the main text. KCl, potassium chloride; AngII, angiotensin II; PE, phenylephrine; ACh, acetylcholine; L-NAME, $N_{\omega}$-nitro-L-arginine methyl ester.

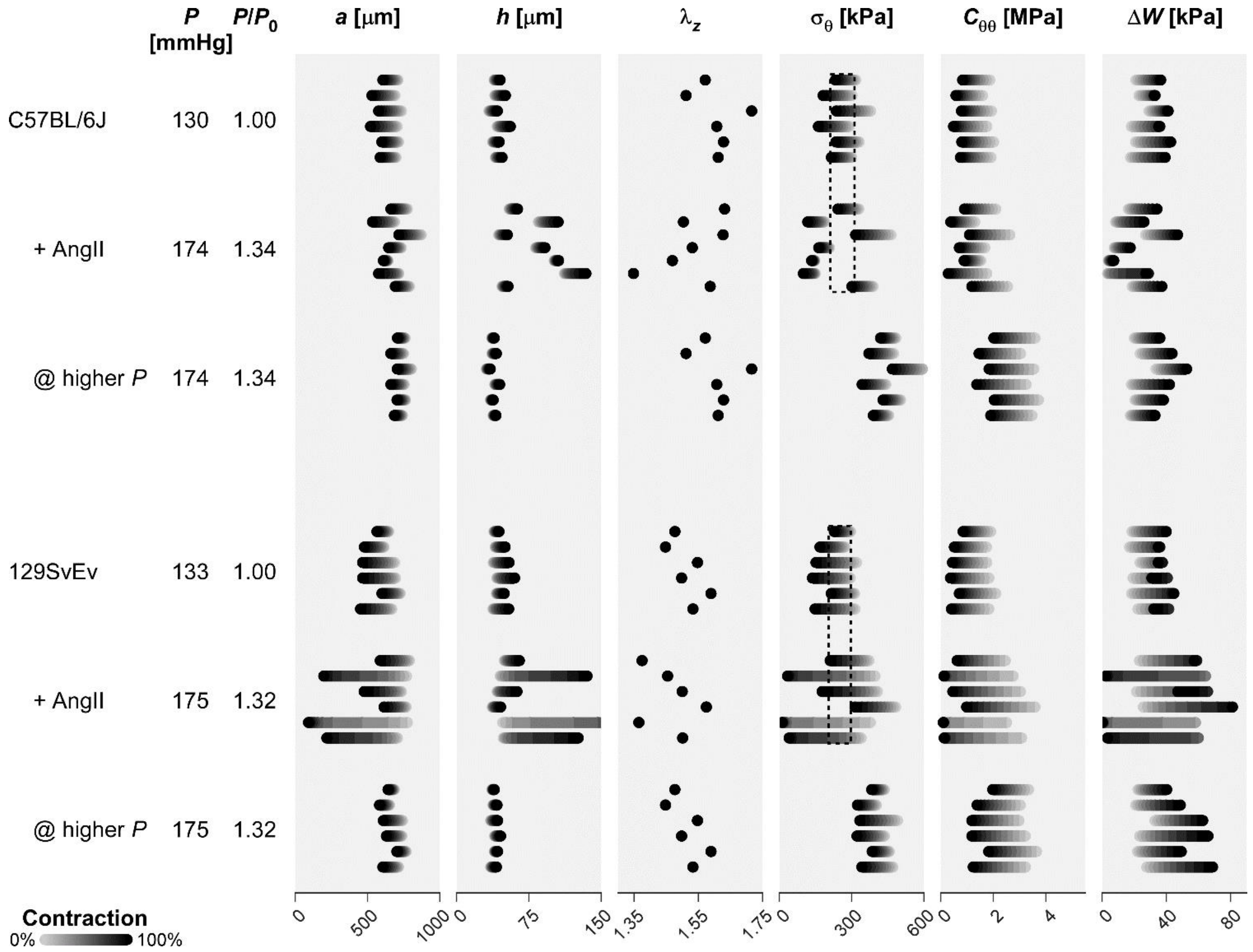

**Supplemental Figure S3.** Similar to Figure S1c except showing the possible effect of smooth muscle cell contraction on values of key biomechanical metrics: contraction ranges from 0% (passive, gray) to 100% (maximally contracted, black). Each horizontally-elongated dot represents one sample; samples are plotted in the order tested. Note that the descending thoracic aorta (DTA) in the C57BL/6J mice is unable to maintain circumferential wall stress at a normotensive level at two-weeks of in vivo AngII infusion. In contrast, except for one sample, the DTA in the 129SvEv mice maintains circumferential wall stress (dotted boxes in $\sigma_\theta$ column) near normal (homeostatic) levels. Finally, note that elastically stored energy is given as the difference between systolic and diastolic states ($\Delta W$) as compared to the absolute stored energy density ($W$) in Figure S1c. This choice was made because smooth muscle contraction shifts the $W = 0$ state (i.e., for a contracted artery, $W \neq 0$ even when traction-free).

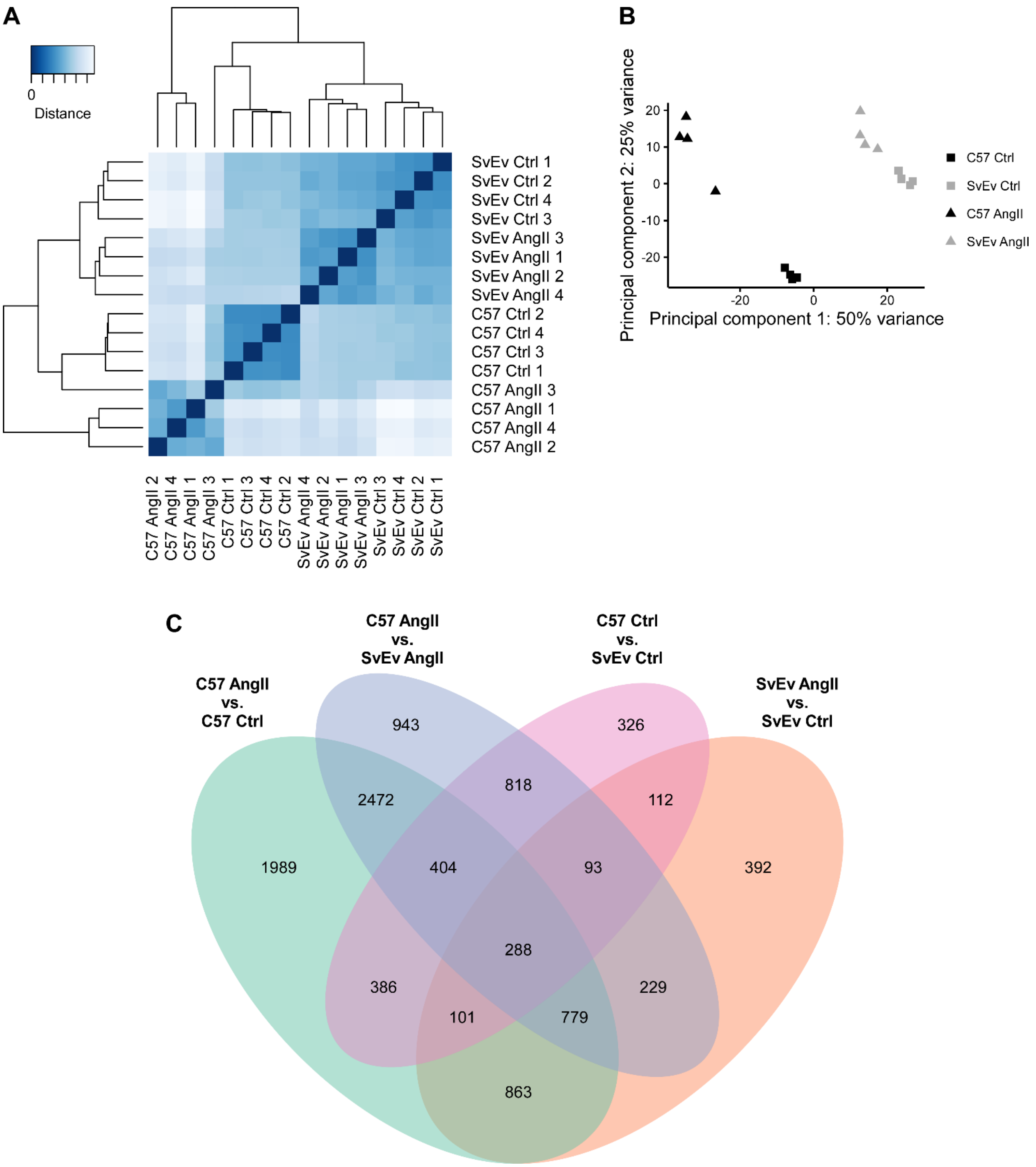

**Supplemental Figure S4. A**: Heatmap of rlog-transformed Euclidean sample distances for the bulk RNAseq data. Samples are numbered 1-4 for each of the four study groups (*n* = 16 total). Despite the fact that ordering was determined automatically based on distances, experimental groups cluster together, confirming that variation in gene expression between groups is larger than within groups, as expected. This finding was also confirmed by a principal component analysis (**B**) of rlog-transformed count data, where samples of the same experimental group clearly cluster together. In addition, qualitatively, AngII infusion causes a bigger difference in gene expression in C57BL/6J (C57) than in 129SvEv (SvEv) mice. **C**: Venn diagram for all significantly differentially expressed genes for all four comparisons. AngII, angiotensin II.

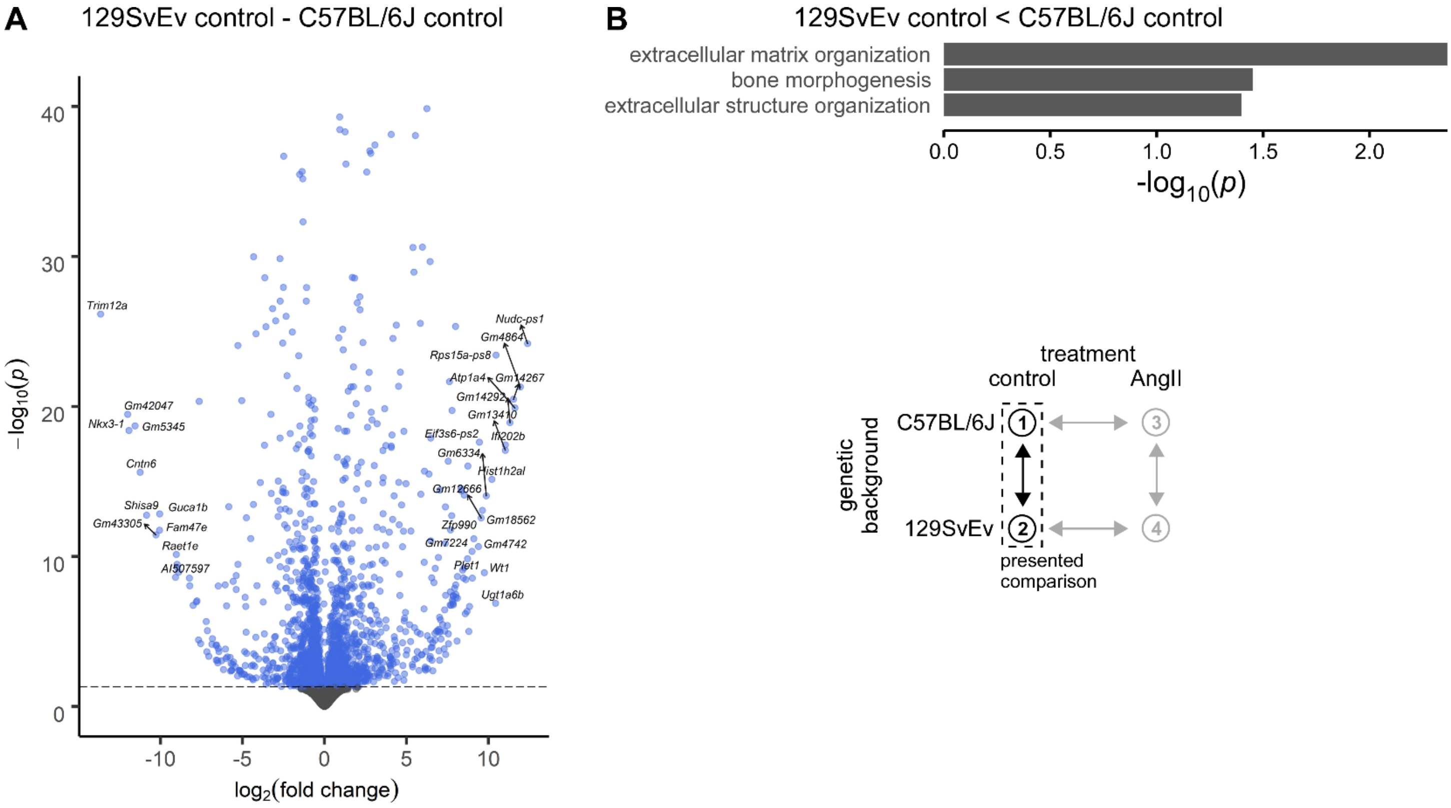

**Supplemental Figure S5a. A**: Volcano plot comparing C57BL/6J and 129SvEv control groups (without angiotensin II (AngII) infusion). y-axis cut off at $-\log_{10}(p)=40$ to avoid compression of volcano base. Dashed line indicates $p=0.05$. **B**: Gene ontology (GO) analysis comparing the same groups to identify differential biological processes. Three ontologies showed significantly higher expression in C57BL/6J than in 129SvEv mice (shown); none showed higher expression in 129SvEv than in C57BL/6J. Note that $-\log_{10}(p)=1.3$ corresponds to $p=0.05$. *p*, Benjamini-Hochberg-adjusted *p*-value.

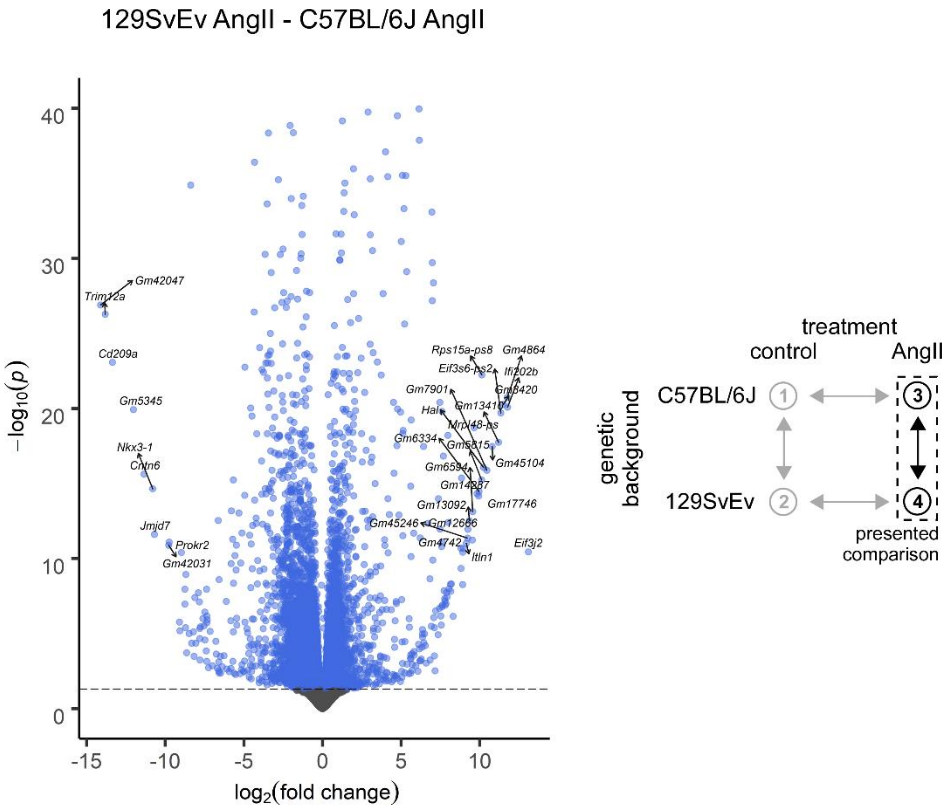

**Supplemental Figure S5b.** Volcano plot comparing angiotensin II (AngII)-infused C57BL/6J and 129SvEv groups. y-axis cut off at $-\log_{10}(p)=40$ to avoid compression of volcano base. *p*, Benjamini-Hochberg-adjusted *p*-value. Dashed line indicates $p=0.05$. See Figure 3 in the main text for the gene ontology (GO) analysis leading to the top-rank biological processes.

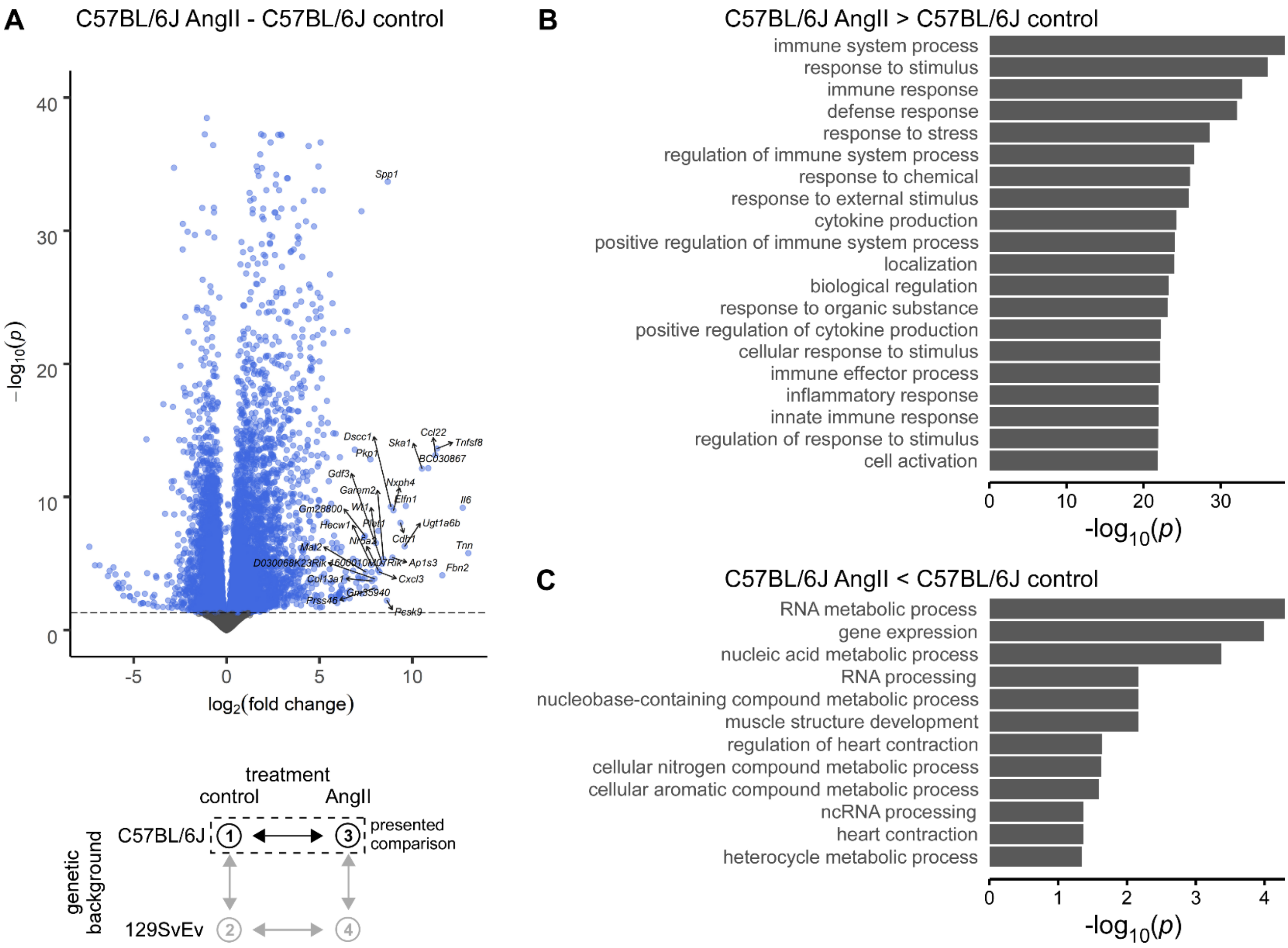

**Supplemental Figure S5c. A**: Volcano plot comparing C57BL/6J groups with and without angiotensin II (AngII) infusion. y-axis cut off at $-\log_{10}(p)=40$ to avoid compression of volcano base. Dashed line indicates $p=0.05$. **B** and **C**: Gene ontology (GO) analysis comparing the same groups. Panel **B** shows the top-20 biological processes (ontologies) that were overexpressed in AngII compared to control; panel **C** shows the ontologies that were under-expressed. Note that $-\log_{10}(p)=1.3$ corresponds to $p=0.05$. *p*, Benjamini-Hochberg-adjusted *p*-value.

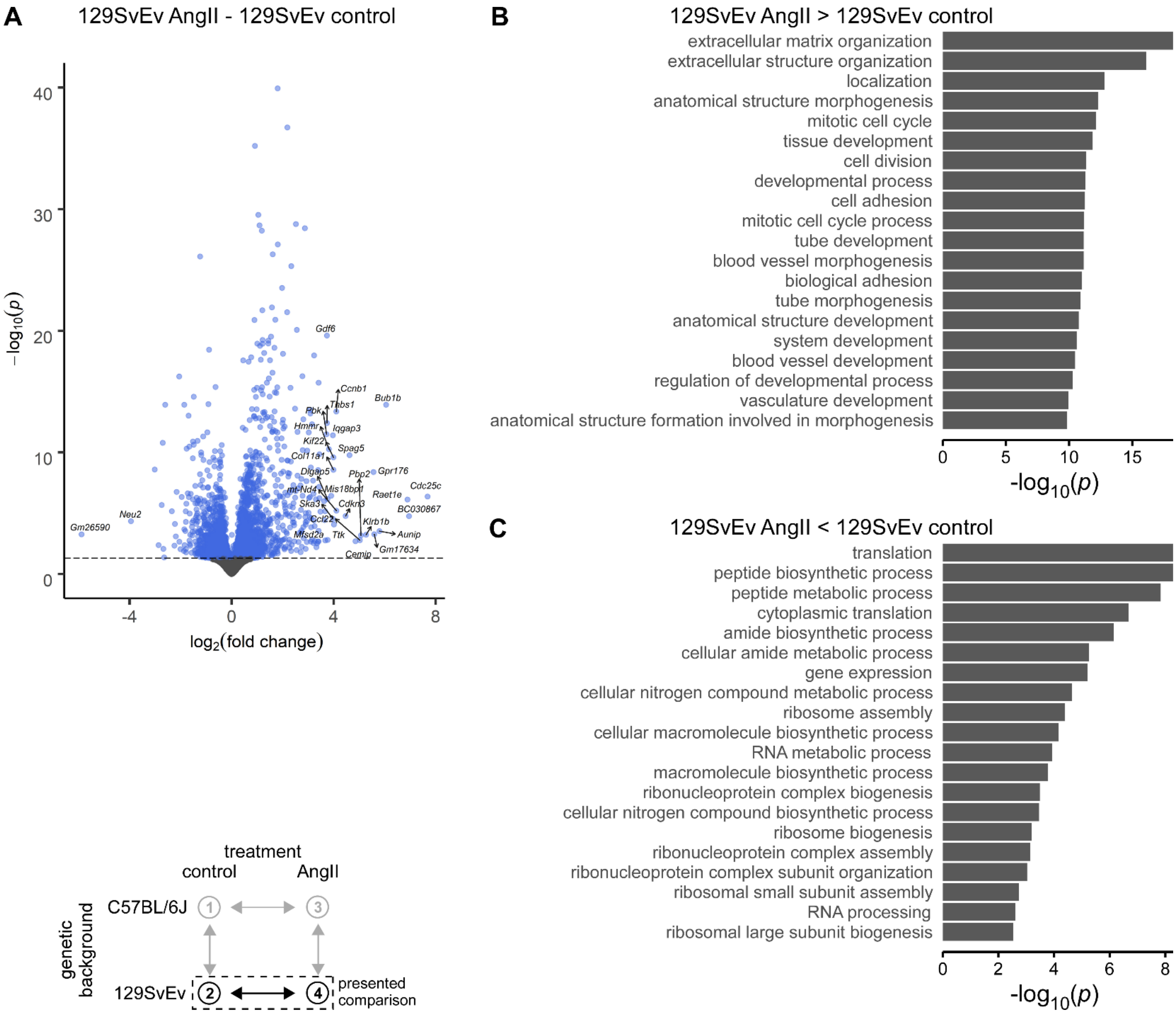

**Supplemental Figure S5d. A**: Volcano plot comparing 129SvEv groups with and without angiotensin II (AngII) infusion. y-axis cut off at -$\log_{10}(p)$=40 to avoid compression of volcano base. Dashed line indicates $p$=0.05. **B** and **C**: Gene ontology (GO) analysis comparing the same groups. Panel **B** shows the top-20 biological processes (ontologies) that were overexpressed in AngII compared to control; note that the following contractility-related ontologies (cf. Figure 3D, main text) were also overexpressed (but not in the top-20): muscle structure development, actin filament-based process, muscle cell differentiation, and actin cytoskeleton organization. Panel **C** shows the top-20 ontologies that were under-expressed in AngII compared to control. Note that -$\log_{10}(p)$=1.3 corresponds to $p$=0.05. $p$, Benjamini-Hochberg-adjusted $p$-value.

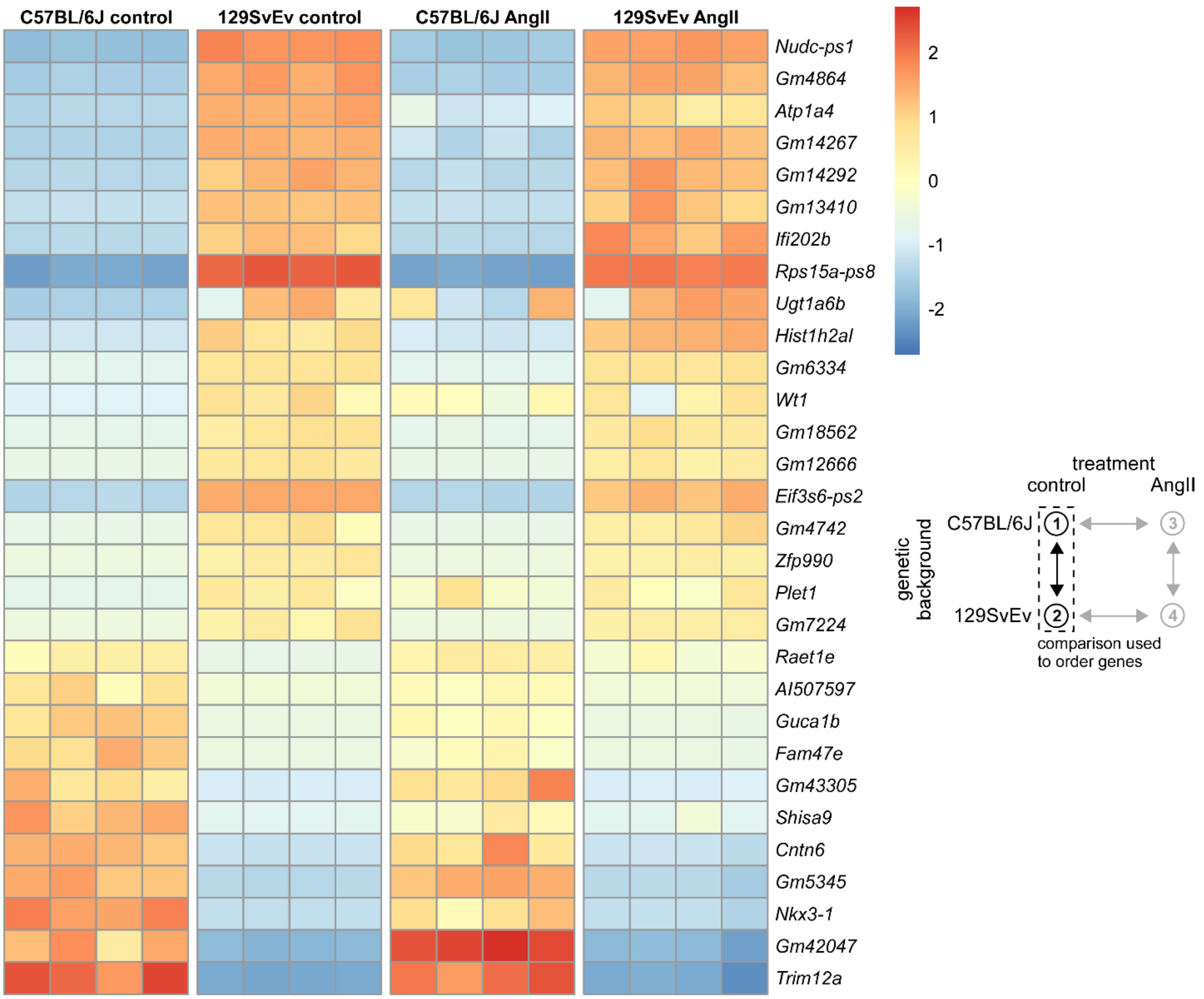

**Supplemental Figure S6a.** rlog-transformed gene expression for all 16 individual samples across the four primary study groups for the top-30 differentially expressed genes that emerged in comparisons between control 129SvEv and C57BL/6J groups. For each gene, mean rlog-transformed expression over all four groups was subtracted for normalization. Ctrl, control; AngII, angiotensin II infused.

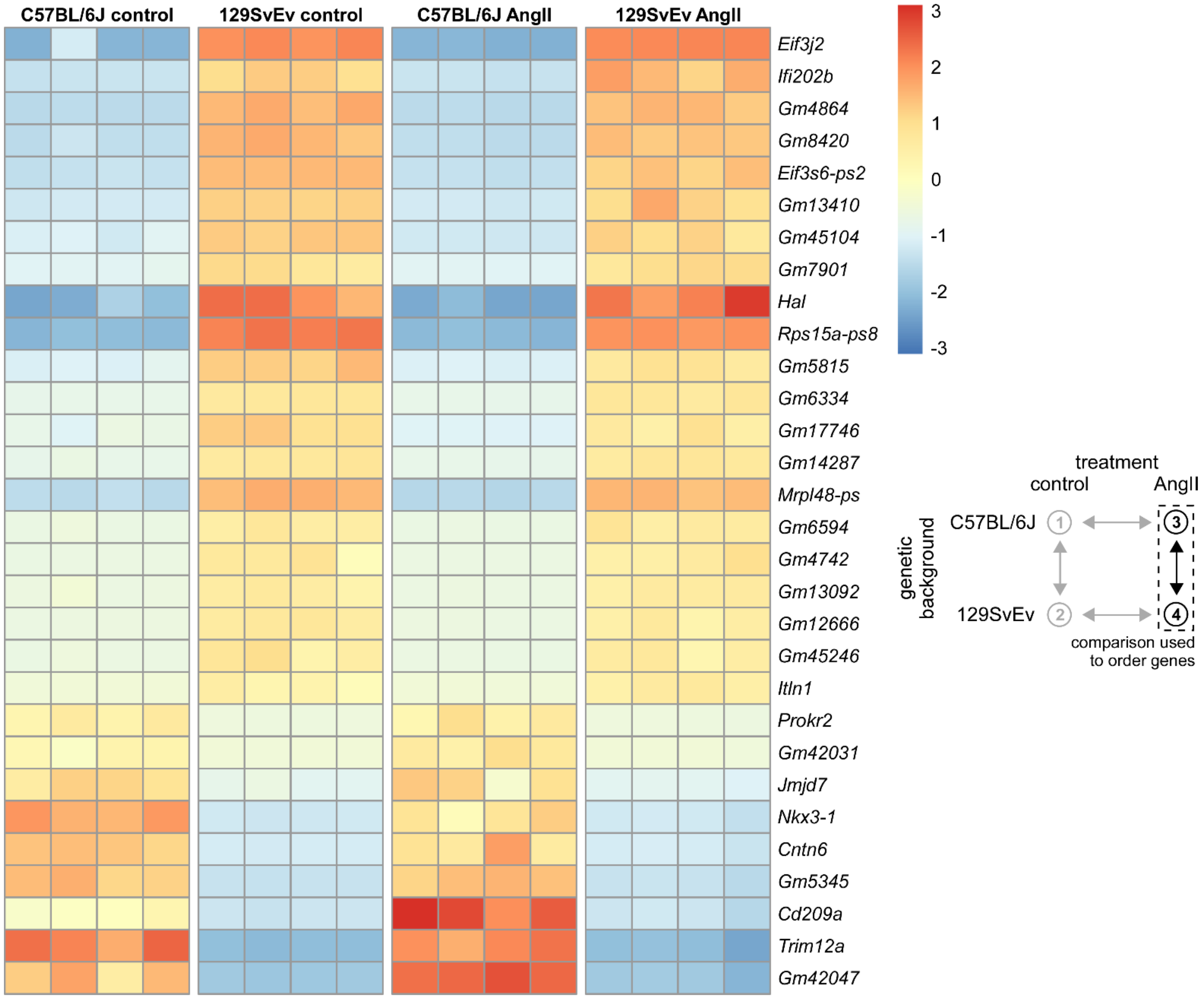

**Supplemental Figure S6b.** rlog-transformed gene expression for all 16 individual samples across the four primary study groups for the top-30 differentially expressed genes that emerged from comparisons between the AngII-infused 129SvEv and C57BL/6J groups. For each gene, mean rlog-transformed expression over all four groups was subtracted for normalization. AngII, angiotensin II infused.

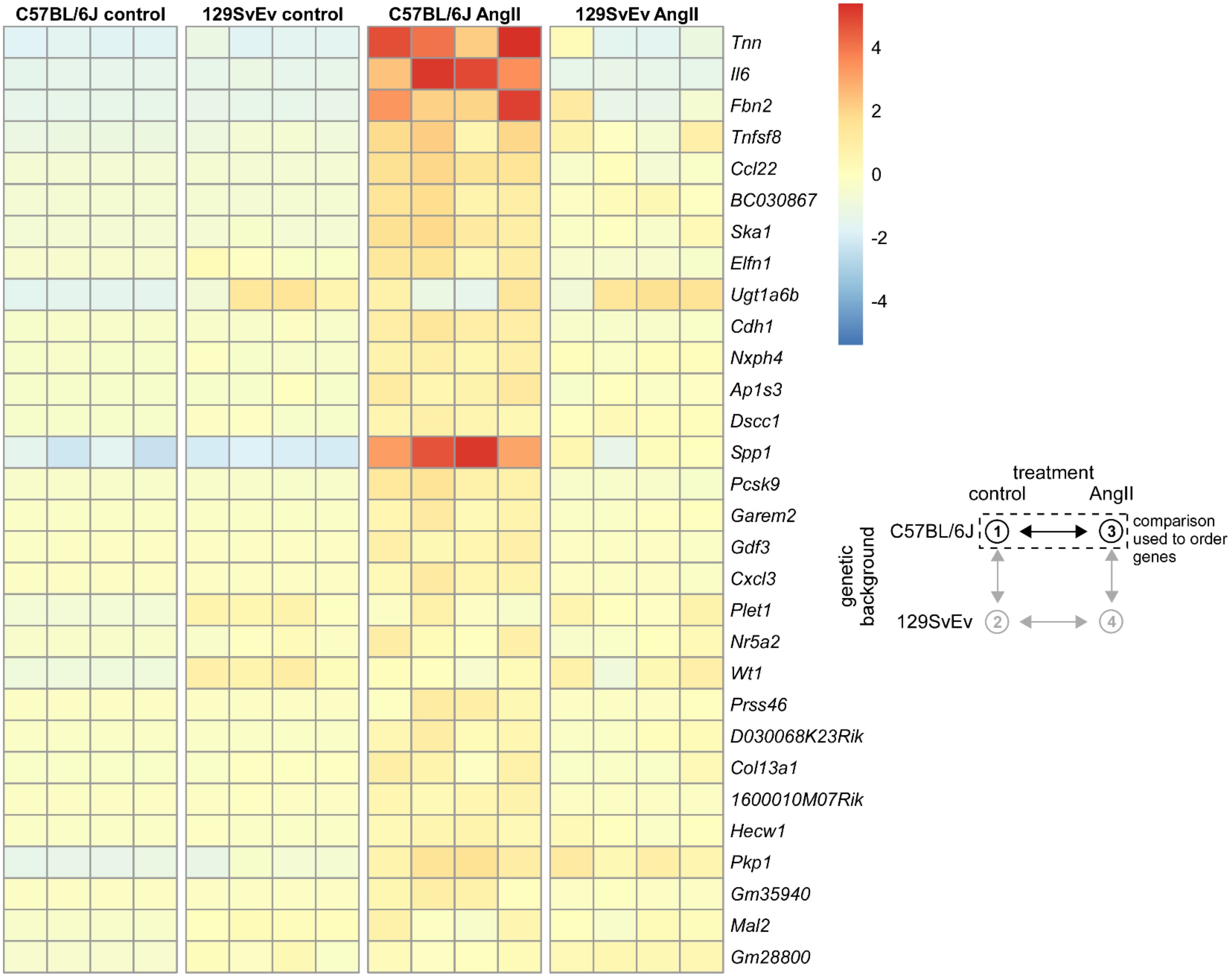

**Supplemental Figure S6c.** rlog-transformed gene expression for all 16 individual samples across the four primary study groups for the top-30 differentially expressed genes that emerged when comparing between the AngII-infused C57BL/6J and control C57BL/6J groups. For each gene, mean rlog-transformed expression over all four groups was subtracted for normalization. AngII, angiotensin II infused.

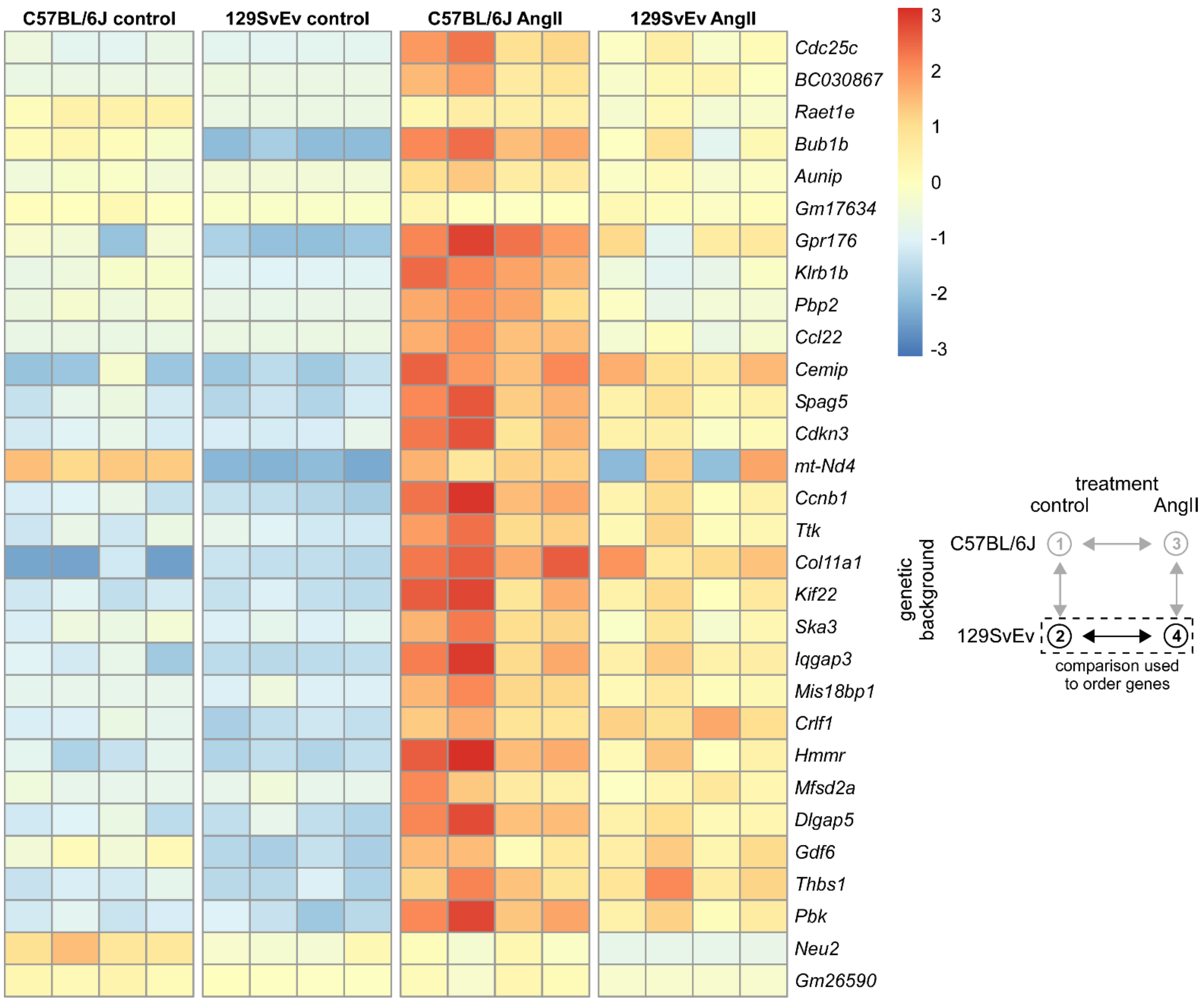

**Supplemental Figure S6d.** rlog-transformed gene expression for all 16 individual samples across the four primary study groups showing the top-30 differentially expressed genes that emerged when comparing between the AngII-infused 129SvEv and control 129SvEv groups. For each gene, mean rlog-transformed expression over all four groups was subtracted for normalization. AngII, angiotensin II infused.

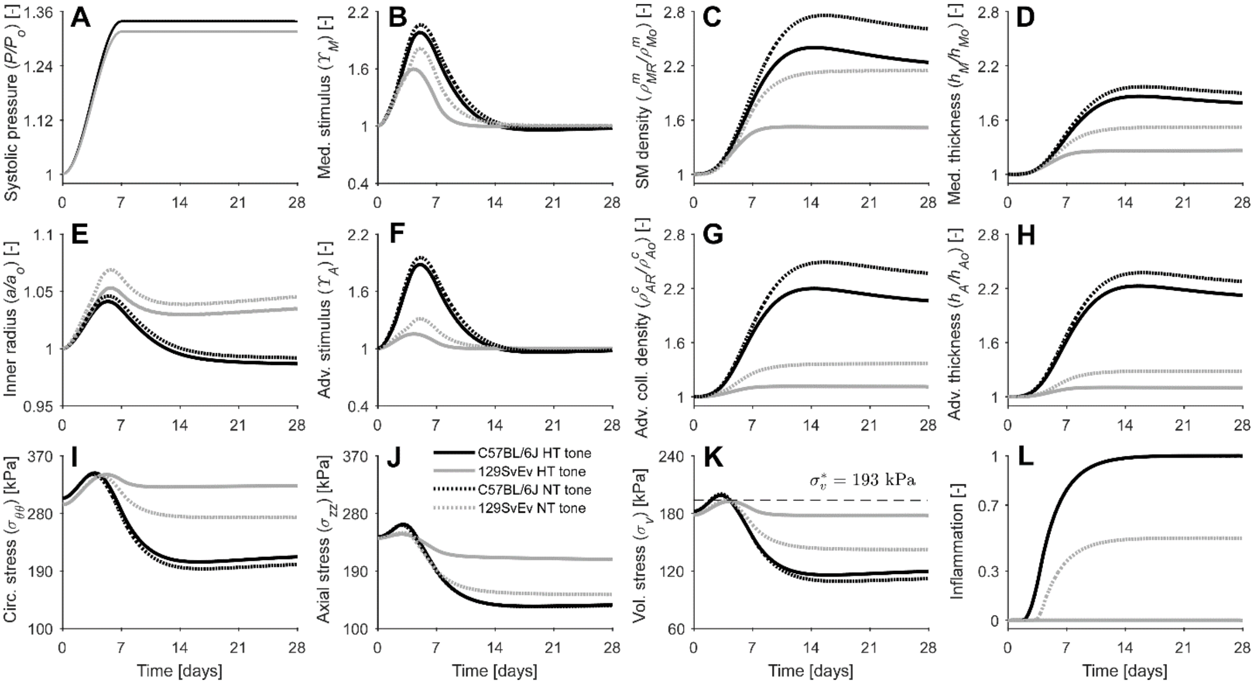

**Figure S7**. Growth & remodeling model predictions of luminal radius (**E**), medial (med., **D**) and adventitial (adv., **H**) thickness (with quantities normalized to values at time 0 prior to AngII infusion) as well as mean circumferential (circ., **I**), axial (**J**), and resulting volumetric (vol., **K**) stress for the C57BL/6J (black) and 129SvEv (grey) mice following respective rapid 1.34- or 1.32-fold increases in systolic pressure (**A**, model input). Shown, too, are the mechano- and inmuno-biological stimulus functions for medial smooth muscle (SM, **B**) and adventitial collagen (coll., **F**) that stimulate respective increases in mass densities (**C** and **G**, normalized). Predictions are shown for a simultaneous increase in smooth muscle tone from day 0 (passive behavior) to respective AngII contraction values under hypertensive (solid lines, AngII-appropriate tone) or normotensive (dotted lines, reduced tone) conditions at day 7, then preserved. Stress-mediated inflammatory effects are computed internally by the model based on a threshold for the volumetric stress (**K**) that determines the onset and extent of the inflammatory response, resulting into marked increases in inflammation for the C57BL/6J mice with both reduced and increased contractility, a moderate increase for the 129SvEv mice with reduced contractility, and a negligible change for the 129SvEv mice with increased contractility (**L**). Note the protective role that the increased vascular tone plays for the 129SvEv mice, by off-loading the early increase in wall stress through contraction and minimizing their inflammatory response, but not for the C57BL/6J mice due to their reduced contractile capacity. AngII, angiotensin II. Compare with Figure 6 in the main text.

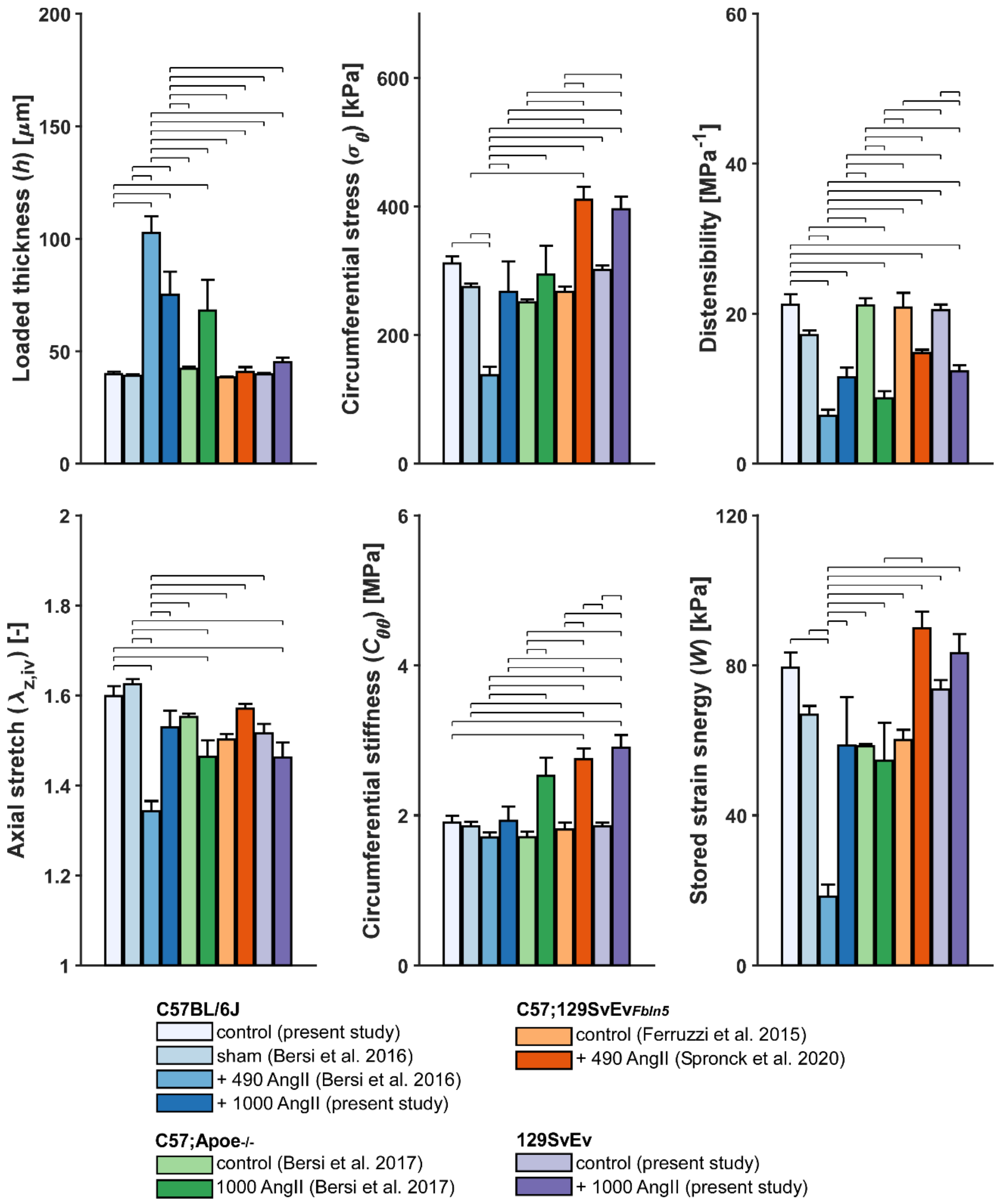

**Supplemental Figure S8.** Effects of angiotensin II (AngII) infusion on select biomechanical metrics (mean ± standard error) across different mouse models, including historical data, all for the descending thoracic aorta (DTA) of adult male mice. AngII doses are given in ng/kg/min. Thin horizontal over-bars denote statistically significant differences ($p$<0.05, one-way analysis of variance with post-hoc Bonferroni tests). Data are from the present as well as three previous studies: Bersi et al. 2016[5] (C57BL/6J), Spronck et al. 2020[7] (C57;129SvEv$_{Fbln5}$), and Bersi et al. 2017[6] (C57;*Apoe*$^{-/-}$). The C57;129SvEv$_{Fbln5}$ mice were generated by breeding *Fbln5*$^{+/-}$ mice to obtain *Fbln5*$^{+/+}$ and *Fbln5*$^{-/-}$ mice. AngII-induced fold changes are shown in Figure S1 and Table S7.

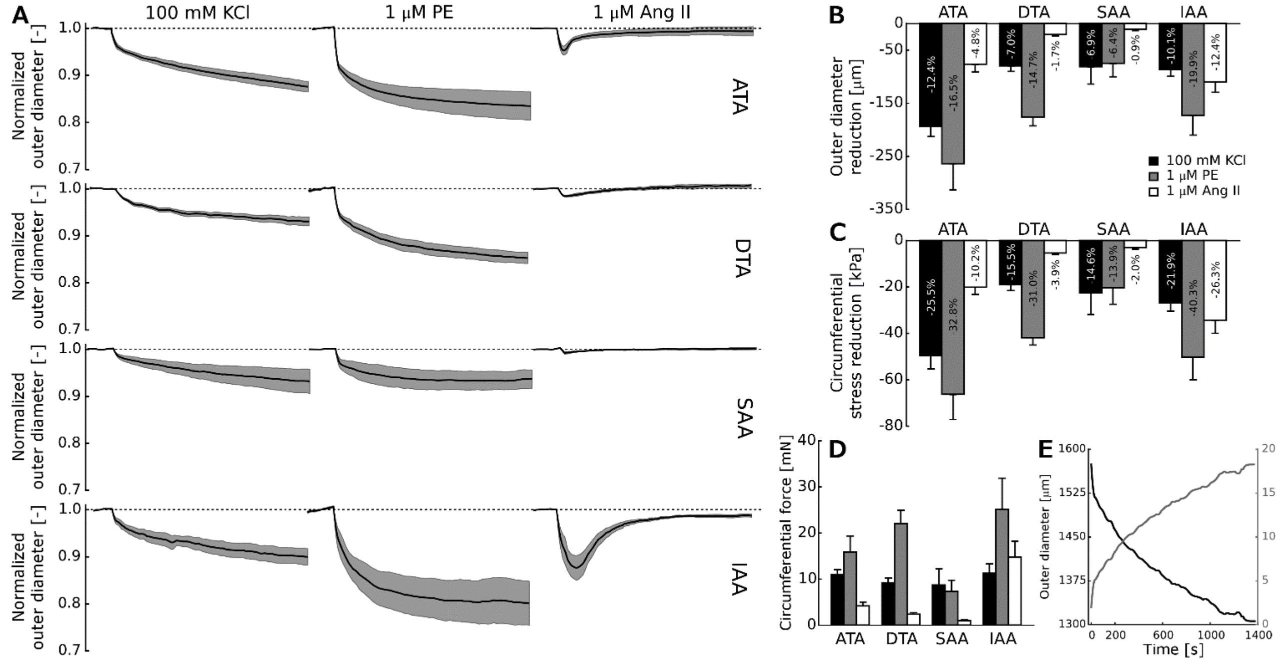

**Supplemental Figure S9. A**: Time-course of vasoconstriction (normalized) induced ex vivo in four regions of the aorta (ATA – ascending thoracic aorta, DTA – descending thoracic aorta, SAA – suprarenal abdominal aorta, and IAA – infrarenal abdominal aorta) from adult male *Apoe*$^{-/-}$ mice on a C57BL/6 background in response to three vaso-stimulants: high potassium chloride (KCl), phenylephrine (PE), and angiotensin (AngII). Note that the response to AngII is significantly greater in the IAA, a region that was previously found to mechano-adapt without fibrosis when hypertension was induced using chronic AngII infusion for 28 days.[6] Shown, too, are the mean maximal reduction in diameter (**B**) and circumferential Cauchy wall stress (**C**) for the four different regions and three vaso-stimulants, again highlighting the more marked response of the IAA to AngII. Finally, show too is an equivalent force generation (as would be measured in ring myography) for the same regions and stimulants, plotted as maximal values (**D**) and time-courses (**E**).

### Supplemental Tables

**Supplemental Table S1.** Passive biomechanical metrics for the descending thoracic aorta harvested from adult male C57BL/6 and 129SvEv mice subjected to normotensive (NT) or AngII-induced hypertensive (HT) conditions.

| | | | | | ANOVA *p*-values | | |
|---|---|---|---|---|---|---|---|
| **Group Number** (for statistics) | **NT C57BL/6J (1)** *n* = 8 | **NT 129SvEv (2)** *n* = 6 | **HT C57BL/6J (3)** *n* = 7 | **HT 129SvEv (4)** *n* = 6 | **strain** | **induced HT** | **strain * induced HT** |
| **Unloaded dimensions** | | | | | | | |
| Wall Thickness, *H* [μm] | 118 ± 2.1 | 110 ± 2.3 | 207 ± 16.0(1)(2) | 130 ± 4.9(3) | **0.000** | **0.000** | **0.001** |
| Inner Diameter, 2*A* [μm] | 664 ± 12 | 649 ± 21 | 625 ± 19 | 659 ± 15 | 0.576 | 0.389 | 0.148 |
| **In Vivo (Loaded) Axial Stretch, $\lambda_z$ [-]** | 1.60 ± 0.02 | 1.52 ± 0.02 | 1.53 ± 0.04 | 1.46 ± 0.03(1) | **0.019** | **0.048** | 0.787 |
| **Dimensions at Diastolic Pressure *P*** [mmHg] | ***P* = 99** | ***P* = 99** | ***P* = 138** | ***P* = 128** | | | |
| Wall Thickness, *h* [μm] | 43 ± 1.1 | 43 ± 0.7 | 78 ± 10.2(1)(2) | 48 ± 2.1(3) | **0.012** | **0.001** | **0.012** |
| Inner Diameter, 2*a* [μm] | 1305 ± 22.2 | 1231 ± 29.9 | 1411 ± 47.6(2) | 1406 ± 17.2(2) | 0.228 | **0.000** | 0.293 |
| **Dimensions at Systolic Pressure *P*** [mmHg] | ***P* = 130** | ***P* = 133** | ***P* = 174** | ***P* = 175** | | | |
| Wall Thickness, *h* [μm] | 40 ± 1.1 | 40 ± 0.7 | 75 ± 10.2(1)(2) | 45 ± 1.9(3) | **0.011** | **0.001** | **0.012** |
| Inner Diameter, 2*a* [μm] | 1419 ± 20.1 | 1345 ± 31.9 | 1490 ± 56.9 | 1514 ± 24.9(2) | 0.506 | **0.003** | 0.194 |
| **Distensibility** [1/MPa] | 21.18 ± 1.38 | 20.44 ± 0.78 | 11.48 ± 1.31(1)(2) | 12.28 ± 0.84(1)(2) | 0.980 | **0.000** | 0.527 |
| **Additional Metrics at Systolic Pressure** | | | | | | | |
| Circumferential Stretch, $\lambda_\theta$ [-] | 1.87 ± 0.03 | 1.83 ± 0.03 | 1.89 ± 0.09 | 1.98 ± 0.05 | 0.659 | 0.113 | 0.262 |
| Circumferential Cauchy Stress, $\sigma_\theta$ [kPa] | 311 ± 11.3 | 301 ± 7.1 | 267 ± 47.6 | 396 ± 19.7(3) | **0.042** | 0.367 | **0.019** |
| Axial Cauchy Stress, $\sigma_z$ [kPa] | 256 ± 13.2 | 250 ± 10.4 | 209 ± 39.9 | 290 ± 16.3 | 0.129 | 0.873 | 0.079 |
| Circumferential Linearized Material Stiffness, $\mathcal{C}_{\theta\theta}$ [MPa] | 1.90 ± 0.09 | 1.85 ± 0.05 | 1.93 ± 0.19 | 2.90 ± 0.17(1)(2)(3) | **0.003** | **0.001** | **0.001** |
| Axial Linearized Material Stiffness, $\mathcal{C}_{zz}$ [MPa] | 2.89 ± 0.11 | 3.61 ± 0.18 | 2.75 ± 0.44 | 4.45 ± 0.40(1)(3) | **0.001** | 0.264 | 0.127 |
| Stored Strain Energy Density, *W* [kPa] | 79 ± 4.1 | 74 ± 2.6 | 59 ± 13.0 | 83 ± 5.1 | 0.232 | 0.480 | 0.059 |
| **Dimensions at Fixed Pressure *P*** [mmHg] | ***P* = 100** | ***P* = 100** | ***P* = 100** | ***P* = 100** | | | |
| Wall Thickness, *h* [μm] | 43 ± 1.1 | 43 ± 0.7 | 85 ± 10.3(1)(2) | 52 ± 2.4(3) | **0.007** | **0.000** | **0.007** |
| Inner Diameter, 2*a* [μm] | 1310 ± 22.1 | 1235 ± 29.9 | 1280 ± 35.5 | 1300 ± 10.7 | 0.322 | 0.523 | 0.090 |
| **Additional Metrics at Fixed Pressure** | ***P* = 100** | ***P* = 100** | ***P* = 100** | ***P* = 100** | | | |
| Circumferential Stretch, $\lambda_\theta$ [-] | 1.73 ± 0.02 | 1.69 ± 0.02 | 1.65 ± 0.05 | 1.72 ± 0.03 | 0.720 | 0.447 | 0.125 |
| Circumferential Cauchy Stress, $\sigma_\theta$ [kPa] | 205 ± 6.5 | 192 ± 4.1 | 112 ± 17.0(1)(2) | 169 ± 8.8(3) | 0.052 | **0.000** | **0.004** |
| Axial Cauchy Stress, $\sigma_z$ [kPa] | 192 ± 10.1 | 197 ± 9.8 | 115 ± 19.0(1)(2) | 171 ± 13.9 | **0.039** | **0.001** | 0.077 |
| Circumferential Linearized Material Stiffness, $\mathcal{C}_{\theta\theta}$ [MPa] | 1.06 ± 0.05 | 0.99 ± 0.02 | 0.59 ± 0.06(1)(2) | 0.86 ± 0.06(3) | 0.061 | **0.000** | **0.003** |
| Axial Linearized Material Stiffness, $\mathcal{C}_{zz}$ [MPa] | 1.79 ± 0.09 | 2.52 ± 0.17(1) | 1.15 ± 0.14(1)(2) | 2.29 ± 0.23(3) | **0.000** | **0.009** | 0.191 |
| Stored Strain Energy Density, *W* [kPa] | 60 ± 2.6 | 54 ± 2.2 | 32 ± 6.2(1)(2) | 46 ± 3.3 | 0.323 | **0.000** | **0.021** |

Values denote mean±standard error. Superscripted numbers in brackets indicate significant differences (*p*<0.05, two-way analysis of variance (ANOVA) followed by Bonferroni post-hoc test), and refer to group numbers.

**Supplemental Table S2.** Active biomechanical metrics for the descending thoracic aorta harvested from adult male C57BL/6 and 129SvEv mice subjected to normotensive (NT) or AngII-induced hypertensive (HT) conditions.

| | **Group Number** (for statistics) | **NT C57BL/6J (1)** ***n*** **= 6** | **NT 129SvEv (2)** ***n*** **= 6** | **HT C57BL/6J (3)** ***n*** **= 7** | **HT 129SvEv (4)** ***n*** **= 6** | **ANOVA *p*-values** strain | induced HT | strain * induced HT |
|---|---|---|---|---|---|---|---|---|
| | **Fixed loading and configuration** | | | | | | | |
| | Axial stretch [-] | 1.49 ± 0.03 | 1.45 ± 0.01 | 1.42 ± 0.04 | 1.41 ± 0.04 | 0.362 | 0.125 | 0.652 |
| | Loaded pressure [mmHg] | 90 | 90 | 90 | 90 | | | |
| | Passive outer diameter @ testing conditions [µm] | 1359 ± 11 | 1292 ± 32 | 1433 ± 27[2] | 1366 ± 11 | **0.008** | **0.004** | 0.989 |
| **100 mM KCl** | **Loaded configuration** | | | | | | | |
| | Relaxed Outer Diameter [µm] | 1297 ± 16.0 | 1239 ± 32.0 | 1339 ± 38.0 | 1295 ± 36.6 | 0.134 | 0.146 | 0.827 |
| | Contracted Outer Diameter [µm] | 1051 ± 26.4 | 958 ± 30.9 | 1059 ± 48.4 | 902 ± 25.5[3] | **0.002** | 0.513 | 0.382 |
| | Change in Outer Diameter [%] | -19 ± 2.1 | -23 ± 0.9 | -21 ± 2.2 | -30 ± 1.4[1,2,3] | **0.001** | **0.013** | 0.140 |
| | **Stress calculations** | | | | | | | |
| | Relaxed Circumferential Stretch [-] | 1.62 ± 0.04 | 1.57 ± 0.02 | 1.49 ± 0.05 | 1.57 ± 0.06 | 0.772 | 0.191 | 0.198 |
| | Contracted Circumferential Stretch [-] | 1.28 ± 0.04 | 1.19 ± 0.01 | 1.12 ± 0.05 | 1.03 ± 0.04[1] | **0.036** | **0.001** | 0.879 |
| | Relaxed Circumferential Stress [kPa] | 149 ± 5.9 | 142 ± 3.2 | 75 ± 10.9[1,2] | 120 ± 6.3[3] | **0.021** | **0.000** | **0.003** |
| | Contracted Circumferential Stress [kPa] | 92 ± 6.3 | 77 ± 2.7 | 39 ± 5.7[1,2] | 49 ± 3.4[1,2] | 0.632 | **0.000** | **0.020** |
| | Change in Circumferential Stress [%] | -38 ± 3.9 | -46 ± 1.5 | -48 ± 4.0 | -59 ± 2.1[1,2] | **0.007** | **0.001** | 0.506 |
| | Active Circumferential Force [mN] | 28 ± 3.4 | 30 ± 2.3 | 34 ± 2.7 | 52 ± 7.1[1,2,3] | **0.028** | **0.005** | 0.078 |
| **1 µM AngII** | **Loaded configuration** | | | | | | | |
| | Relaxed Outer Diameter [µm] | 1300 ± 13.8 | 1232 ± 29.1 | 1348 ± 33.9 | 1280 ± 47.1 | 0.055 | 0.167 | 0.993 |
| | Contracted Outer Diameter [µm] | 1251 ± 23.0 | 1094 ± 34.8[1] | 1215 ± 40.0 | 995 ± 42.8[1,3] | **0.000** | 0.078 | 0.396 |
| | Change in Outer Diameter [%] | -4 ± 1.2 | -11 ± 2.0[1] | -10 ± 1.5 | -22 ± 1.2[1,2,3] | **0.000** | **0.000** | 0.114 |
| | **Stress calculations** | | | | | | | |
| | Relaxed Circumferential Stretch [-] | 1.62 ± 0.03 | 1.56 ± 0.02 | 1.51 ± 0.05 | 1.55 ± 0.07 | 0.848 | 0.201 | 0.299 |
| | Contracted Circumferential Stretch [-] | 1.56 ± 0.04 | 1.38 ± 0.02 | 1.35 ± 0.06[1] | 1.19 ± 0.06[1] | **0.003** | **0.001** | 0.842 |
| | Relaxed Circumferential Stress [kPa] | 150 ± 4.9 | 140 ± 2.5 | 76 ± 10.6[1,2] | 117 ± 8.7[3] | 0.056 | **0.000** | **0.004** |
| | Contracted Circumferential Stress [kPa] | 137 ± 4.6 | 107 ± 5.8 | 58 ± 9.3[1,2] | 64 ± 7.2[1,2] | 0.104 | **0.000** | **0.022** |
| | Change in Circumferential Stress [%] | -8 ± 2.6 | -24 ± 4.1[1] | -24 ± 3.5[1] | -46 ± 2.5[1,2,3] | **0.000** | **0.000** | 0.289 |
| | Active Circumferential Force [mN] | 5 ± 2.0 | 14 ± 3.0 | 14 ± 2.0 | 34 ± 4.0[1,2,3] | **0.000** | **0.000** | 0.055 |
| **1 µM PE** | **Loaded configuration** | | | | | | | |
| | Relaxed Outer Diameter [µm] | 1330 ± 12.1 | 1252 ± 32.7 | 1383 ± 29.8[2] | 1306 ± 33.7 | **0.014** | 0.077 | 0.969 |
| | Contracted Outer Diameter [µm] | 1008 ± 25.5 | 937 ± 32.7 | 974 ± 35.1 | 800 ± 38.9[1,3] | **0.002** | **0.019** | 0.144 |
| | Change in Outer Diameter [%] | -24 ± 1.8 | -25 ± 2.8 | -30 ± 2.2 | -39 ± 2.5[1,3] | **0.046** | **0.001** | 0.087 |
| | **Stress calculations** | | | | | | | |
| | Relaxed Circumferential Stretch [-] | 1.66 ± 0.03 | 1.59 ± 0.02 | 1.55 ± 0.05 | 1.58 ± 0.05 | 0.619 | 0.185 | 0.221 |
| | Contracted Circumferential Stretch [-] | 1.22 ± 0.04 | 1.15 ± 0.04 | 0.99 ± 0.05[1] | 0.87 ± 0.05[1,2] | 0.056 | **0.000** | 0.636 |
| | Relaxed Circumferential Stress [kPa] | 158 ± 5.8 | 145 ± 2.9 | 83 ± 13.0[1,2] | 123 ± 7.5[3] | 0.126 | **0.000** | **0.007** |
| | Contracted Circumferential Stress [kPa] | 83 ± 5.2 | 74 ± 6.0 | 30 ± 5.4[1,2] | 35 ± 7.4[1,2] | 0.769 | **0.000** | 0.252 |
| | Change in Circumferential Stress [%] | -47 ± 3.2 | -49 ± 4.6 | -64 ± 3.5[1] | -72 ± 4.1[1,2] | 0.205 | **0.000** | 0.373 |
| | Active Circumferential Force [mN] | 39 ± 4.1 | 35 ± 5.2 | 56 ± 5.4 | 76 ± 10.1[1,2] | 0.224 | **0.000** | 0.076 |

Table continues on next page.

| | | | | | | | |
|---|---|---|---|---|---|---|---|
| **10 µM ACh / 1 mM L-NAME** | | | | | | | |
| **Loaded configuration** | | | | | | | |
| Relaxed Outer Diameter w ACh [µm] | 1196 ± 20.0 | 1073 ± 41.1 | 1170 ± 46.2 | 918 ± 62.2(1)(3) | **0.000** | 0.060 | 0.170 |
| Contracted Outer Diameter w L-NAME [µm] | 867 ± 22.4 | 780 ± 32.1 | 887 ± 39.0 | 665 ± 22.8(1)(3) | **0.000** | 0.140 | **0.041** |
| Max EC activation [µm] (L-NAME od - ACh od) | 329 ± 29.8 | 292 ± 22.9 | 283 ± 46.3 | 253 ± 61.8 | 0.453 | 0.339 | 0.942 |
| Basal EC activation [µm] (ΔL-NAME – ΔPE) | 141 ± 15.3 | 157 ± 22.7 | 86 ± 20.9 | 135 ± 31.4 | 0.174 | 0.114 | 0.487 |
| Max SMC activation [µm] (passive - L-NAME od) | -492 ± 23.5 | -512 ± 34.2 | -545 ± 41.7 | -706 ± 20.6(1)(2)(3) | **0.011** | **0.001** | **0.041** |
| Basal SMC activation [µm] (tone - basal EC) | -170 ± 12.1 | -198 ± 20.5 | -136 ± 15.8 | -200 ± 28.6 | **0.033** | 0.437 | 0.376 |
| Total basal tone [µm] (basal PE - passive od) | -29 ± 6.3 | -41 ± 5.2 | -50 ± 11.4 | -65 ± 23.2 | 0.346 | 0.110 | 0.903 |
| **Stress calculations** | | | | | | | |
| Relaxed Circ Stress w ACh [kPa] | 123.87 ± 4.81 | 102.98 ± 9.30 | 54.60 ± 10.95(1)(2) | 52.19 ± 9.50(1)(2) | 0.219 | **0.000** | 0.326 |
| Contracted Circ Stress w L-NAME [kPa] | 56.52 ± 4.53 | 45.30 ± 4.94 | 20.56 ± 3.17(1)(2) | 17.34 ± 4.04(1)(2) | 0.097 | **0.000** | 0.347 |
| Max EC activation [kPa] (L-NAME - ACh) | 67 ± 5.3 | 58 ± 6.0 | 34 ± 8.4(1) | 35 ± 9.0(1) | 0.561 | **0.001** | 0.493 |
| Basal EC activation [kPa] (ΔL-NAME - ΔPE) | 26 ± 2.7 | 29 ± 4.6 | 9 ± 3.1(1)(2) | 18 ± 4.8 | 0.179 | **0.002** | 0.429 |
| Max SMC activation [kPa] (passive - L-NAME) | -109 ± 5.0 | -111 ± 5.8 | -69 ± 11.3(1)(2) | -120 ± 4.6(3) | **0.003** | 0.062 | **0.004** |
| Basal SMC activation [kPa] (tone - basal EC) | -34 ± 2.1 | -39 ± 4.3 | -17 ± 2.9(1)(2) | -33 ± 6.1 | **0.016** | **0.007** | 0.202 |
| Total basal tone [kPa] (basal PE - passive od) | -8 ± 1.8 | -11 ± 1.5 | -7 ± 2.0 | -15 ± 5.1 | 0.089 | 0.598 | 0.456 |
| **Metrics normalized to max SMC** | | | | | | | |
| Max EC diameter increase [%] | 66.43 ± 3.7 | 59.14 ± 7.11 | 51.07 ± 6.14 | 35.22 ± 7.89(1) | 0.086 | **0.006** | 0.512 |
| Basal EC diameter increase [%] | 28.83 ± 3.4 | 31.02 ± 4.75 | 15.45 ± 3.24 | 19.06 ± 4.34 | 0.470 | **0.004** | 0.858 |
| Basal SMC diameter decrease [%] | 34.91 ± 3.0 | 39.32 ± 4.5 | 24.74 ± 1.8(2) | 28.45 ± 4.2 | 0.253 | **0.006** | 0.921 |
| Total tone diameter decrease [%] | 6.08 ± 1.4 | 8.30 ± 1.6 | 9.30 ± 2.0 | 9.40 ± 3.3 | 0.603 | 0.336 | 0.634 |
| Max EC circ stress increase [%] | 61.75 ± 3.9 | 53.82 ± 7.65 | 46.28 ± 5.72 | 28.91 ± 7.43(1) | 0.059 | **0.004** | 0.464 |
| Basal EC circ stress increase [%] | 24.51 ± 3.2 | 26.02 ± 4.45 | 12.85 ± 2.69 | 14.53 ± 3.52 | 0.651 | **0.003** | 0.981 |
| Basal SMC circ stress decrease [%] | 31.79 ± 2.9 | 36.07 ± 4.2 | 23.89 ± 1.4 | 26.53 ± 4.2 | 0.301 | **0.014** | 0.803 |
| Total tone circ stress decrease [%] | 7.28 ± 1.6 | 10.05 ± 1.8 | 11.04 ± 2.3 | 12.00 ± 4.2 | 0.491 | 0.295 | 0.736 |

Values denote mean±standard error. Superscripted numbers in brackets indicate significant differences ($p$<0.05, two-way analysis of variance (ANOVA) followed by Bonferroni post-hoc test), and refer to group numbers.

**Supplemental Table S3.** Quantitative histology to determine elastin, smooth muscle, collagen, and glycosaminoglycan contents for the study groups (C57BL/6 and 129SvEv mice) subjected to normotensive (NT) or AngII-induced hypertensive (HT) conditions.

| | **Group Number** (for statistics) | **NT C57BL/6J (1)** | **NT 129SvEv (2)** | **HT C57BL/6J (3)** | **HT 129SvEv (4)** | **ANOVA *p*-values** strain | induced HT | strain* induced HT |
|---|---|---|---|---|---|---|---|---|
| **Unloaded Cross-sectional Area** [mm²] | | | | | | | | |
| Media | Elastin | 0.068 ± 0.002 | 0.079 ± 0.004 | 0.089 ± 0.007 | 0.101 ± 0.014 | 0.203 | **0.027** | 0.915 |
| Media | Smooth muscle | 0.073 ± 0.005 | 0.083 ± 0.007 | 0.165 ± 0.013(1)(2) | 0.122 ± 0.004(1)(2)(3) | 0.075 | **0.000** | **0.006** |
| Media | Collagen | 0.045 ± 0.006 | 0.036 ± 0.005 | 0.080 ± 0.012(2) | 0.021 ± 0.015(3) | **0.005** | 0.380 | **0.031** |
| Media | Glycosaminoglycans | 0.004 ± 0.001 | 0.005 ± 0.001 | 0.007 ± 0.001 | 0.007 ± 0.001 | 0.545 | **0.010** | 0.179 |
| Media | Total | 0.190 ± 0.011 | 0.204 ± 0.007 | 0.341 ± 0.030(1)(2) | 0.251 ± 0.003(3) | **0.042** | **0.000** | **0.007** |
| Adventitia | Elastin | 0.000 ± 0.000 | 0.000 ± 0.000 | 0.001 ± 0.000(1)(2) | 0.001 ± 0.000(3) | 0.056 | **0.005** | 0.056 |
| Adventitia | Smooth muscle | - | - | - | - | | | |
| Adventitia | Collagen | 0.088 ± 0.005 | 0.065 ± 0.003 | 0.206 ± 0.035(1)(2) | 0.069 ± 0.002(3) | **0.000** | **0.004** | **0.007** |
| Adventitia | Glycosaminoglycans | - | - | - | - | | | |
| Adventitia | Total | 0.089 ± 0.005 | 0.065 ± 0.003 | 0.207 ± 0.035(1)(2) | 0.070 ± 0.002(3) | **0.000** | **0.004** | **0.008** |
| Total (both layers) | | 0.280 ± 0.006 | 0.270 ± 0.010 | 0.548 ± 0.025(1)(2) | 0.321 ± 0.005(3) | **0.000** | **0.000** | **0.000** |
| **Loaded Cross-sectional Area** [mm²] | | | | | | | | |
| Media | Elastin | 0.044 ± 0.002 | 0.052 ± 0.003 | 0.057 ± 0.005 | 0.069 ± 0.010 | 0.138 | **0.024** | 0.770 |
| Media | Smooth muscle | 0.047 ± 0.002 | 0.054 ± 0.004 | 0.106 ± 0.009(1)(2) | 0.082 ± 0.003(1)(2)(3) | 0.153 | **0.000** | **0.013** |
| Media | Collagen | 0.029 ± 0.003 | 0.024 ± 0.003 | 0.051 ± 0.008(2) | 0.014 ± 0.010(3) | **0.007** | 0.387 | **0.033** |
| Media | Glycosaminoglycans | 0.002 ± 0.000 | 0.004 ± 0.000 | 0.005 ± 0.001 | 0.004 ± 0.000 | 0.371 | **0.008** | 0.224 |
| Media | Total | 0.122 ± 0.005 | 0.132 ± 0.003 | 0.219 ± 0.020(1)(2) | 0.169 ± 0.003(3) | 0.097 | **0.000** | **0.012** |
| Adventitia | Elastin | 0.000 ± 0.000 | 0.000 ± 0.000 | 0.001 ± 0.000(1)(2) | 0.000 ± 0.000 | 0.071 | **0.005** | 0.072 |
| Adventitia | Smooth muscle | - | - | - | - | | | |
| Adventitia | Collagen | 0.057 ± 0.004 | 0.042 ± 0.002 | 0.134 ± 0.024(1)(2) | 0.047 ± 0.001(3) | **0.001** | **0.005** | **0.011** |
| Adventitia | Glycosaminoglycans | - | - | - | - | | | |
| Adventitia | Total | 0.057 ± 0.004 | 0.042 ± 0.002 | 0.135 ± 0.024(1)(2) | 0.047 ± 0.001(3) | **0.001** | **0.005** | **0.012** |
| Total (both layers) | | 0.179 ± 0.001 | 0.175 ± 0.005 | 0.354 ± 0.020(1)(2) | 0.216 ± 0.004(3) | **0.000** | **0.000** | **0.000** |
| **Percentage of per-layer area** [%] | | | | | | | | |
| Media | Elastin | 36.5 ± 2.2 | 39.5 ± 3.0 | 26.5 ± 1.2 | 40.2 ± 5.3(3) | **0.026** | 0.201 | 0.144 |
| Media | Smooth muscle | 38.4 ± 0.6 | 40.1 ± 2.4 | 48.9 ± 1.5(1)(2) | 48.5 ± 1.2(1)(2) | 0.724 | **0.000** | 0.549 |
| Media | Collagen | 23.2 ± 2.0 | 17.7 ± 2.3 | 22.5 ± 2.3 | 8.7 ± 5.8 | **0.017** | 0.211 | 0.279 |
| Media | Glycosaminoglycans | 1.9 ± 0.2 | 2.7 ± 0.4 | 2.1 ± 0.3 | 2.6 ± 0.3 | **0.048** | 0.934 | 0.773 |
| Adventitia | Elastin | 0.5 ± 0.1 | 0.7 ± 0.1 | 0.7 ± 0.1 | 1.0 ± 0.1 | 0.131 | 0.121 | 0.641 |
| Adventitia | Smooth muscle | - | - | - | - | | | |
| Adventitia | Collagen | 99.5 ± 0.1 | 99.3 ± 0.1 | 99.3 ± 0.1 | 99.0 ± 0.1 | 0.131 | 0.121 | 0.641 |
| Adventitia | Glycosaminoglycans | - | - | - | - | | | |
| **Percentage of total area** [%] | | | | | | | | |
| Media | | 67.8 ± 2.5 | 75.8 ± 0.5 | 63.0 ± 5.5(2) | 78.2 ± 0.4(3) | **0.001** | 0.711 | 0.267 |
| Adventitia | | 32.0 ± 2.4 | 24.2 ± 0.5 | 37.0 ± 5.5(2) | 21.8 ± 0.4(3) | **0.001** | 0.689 | 0.253 |

Values denote mean±standard error. Superscripted numbers in brackets indicate significant differences ($p<0.05$, two-way analysis of variance (ANOVA) followed by Bonferroni post-hoc test), and refer to group numbers.

**Supplemental Table S4.** Quantitative histology to determine CD45-positive area for the C57BL/6 and 129SvEv mice subjected to normotensive (NT) or AngII-induced hypertensive (HT) conditions

| | | | | | ANOVA *p*-values | | |
|---|---|---|---|---|---|---|---|
| **Group Number** (for statistics) | **NT C57BL/6J (1)** | **NT 129SvEv (2)** | **HT C57BL/6J (3)** | **HT 129SvEv (4)** | **strain** | **induced HT** | **strain * induced HT** |
| **CD45-positive area** | | | | | | | |
| Unloaded [µm$^2$] | 348 ± 91 | 3008 ± 413 | 7831 ± 1483$^{(1)(2)}$ | 2614 ± 708$^{(3)}$ | 0.161 | **0.000** | **0.000** |
| Loaded [µm$^2$] | 220 ± 55 | 1971 ± 284 | 5043 ± 955$^{(1)(2)}$ | 1754 ± 470$^{(3)}$ | 0.194 | **0.000** | **0.000** |
| Percentage of total area [%] | 0.123 ± 0.030 | 1.150 ± 0.184$^{(1)}$ | 1.457 ± 0.304$^{(1)}$ | 0.795 ± 0.208 | 0.417 | **0.036** | **0.001** |
| Ratio of area percentage WRT NT [-] | 1.000 ± 0.248 | 1.000 ± 0.160 | 11.884 ± 2.482$^{(1)(2)}$ | 0.691 ± 0.181$^{(3)}$ | **0.000** | **0.000** | **0.000** |

Values denote mean±standard error. Superscripted numbers in brackets indicate significant differences ($p<0.05$, two-way analysis of variance (ANOVA) followed by Bonferroni post-hoc test), and refer to group numbers.

**Supplemental Table S5**. Maximum contractility is a stronger predictor of hypertensive wall thickening across all groups than is $CD45^+$ content.

| | Standardized β | | *F*-test (model comparison) |
|---|---|---|---|
| | **coefficient** | ***p-value*** | ***p*-value** |
| **Model #1** | | | |
| Constant | - | - | |
| CD45+ cross-sectional area | 0.846 | 0.034 | |
| **Model #2** | | | |
| Constant | - | - | |
| Maximum active Cauchy stress reduction | 0.972 | 0.001 | |
| **Model #3** | | | |
| Constant | - | - | vs. #1: 0.025 |
| CD45+ cross-sectional area | 0.195 | 0.393 | vs. #2: 0.393 |
| Maximum active Cauchy stress reduction | 0.816 | 0.025 | |

Multivariable linear regression assessment predicting wall thickness from contractility and/or CD45+ cross-sectional area. Analysis performed on hypertensive data groups only and only samples for which both contractility and $CD45^+$ histology data were available. Wall thicknesses were evaluated at a common, loaded pressure of 100 mmHg and at individual-specific axial stretches.

**Supplemental Table S6.** Comparison of passive biomechanical metrics for descending thoracic aortas (DTAs) of normotensive (NT) mice of four different background strains. Note that these passive baseline mechanical properties of our two pure strains (C57BL/6J and 129SvEv) and two mixed strains (C57BL/6;129SvEv)[2,7] are very similar.

| **Mouse Model (NT)** | **Systolic Pressure (*P*) [mmHg]** | **Inner Diameter (2*a*) [μm]** | **Wall Thickness (*h*) [μm]** | **Axial Stretch Ratio ($\lambda_z$)** | **Circ. Stress ($\sigma_\theta$) [kPa]** | **Circ. Stiffness ($C_{\theta\theta}$) [MPa]** | **Stored Energy (*W*) [kPa]** |
|---|---|---|---|---|---|---|---|
| C57BL/6J (*n*=8)(1) | 130 | 1419 ± 20 | 39.8 ± 1.1 | 1.60±0.02 | 311 ± 11 | 1.90±0.09 | 79 ± 4 |
| C57;129SvEv$_{Fbln5}$ (*n*=5)(2) | 120 | 1277 ± 28(1) | 38.3 ± 0.4 | 1.50±0.01 | 267 ± 8 | 1.81±0.09 | 60 ± 3(1) |
| C57;129SvEv$_{Myh11}$ (*n*=5)(3) | 124 | 1342 ± 38 | 41.5 ± 2.2 | 1.54±0.04 | 269 ± 9 | 2.26±0.1(1)(2) | 57 ± 3(1) |
| 129SvEv (*n*=6)(4) | 133 | 1345 ± 32 | 39.6 ± 0.7 | 1.52±0.02 | 301 ± 7 | 1.85±0.05(3) | 74 ± 3(3) |

Mean±standard error values of systolic pressure (measured using a tail-cuff method) plus the associated inner diameter and wall thickness, in vivo value of axial wall stretch, circumferential wall stress and material stiffness, and elastically stored energy. Superscripted numbers indicate statistically significant differences across the four groups (1–4) using a one-way analysis of variance with post-hoc Bonferroni tests. Note the general similarity across models, consistent with that found previously for the ascending aorta.[6]

**Supplemental Table S7**. Relative changes in passive biomechanical metrics in response to AngII-induced hypertension. All data are from previous studies using the same methods of data collection and analysis and are included for direct comparison with the present data.

| | **Systolic Pressure (Absolute/ Ratio)** | | **Inner Diameter Ratio** | **Wall Thickness Ratio** | **Axial Stretch Ratio** | **Circ. Stress Ratio** | **Circ. Stiffness Ratio** | **Stored Energy Ratio** |
|---|---|---|---|---|---|---|---|---|
| C57BL/6J[1] | 121 | 1.00 | 1.00±0.01 | 1.00±0.02 | 1.00±0.01 | 1.00±0.02 | 1.00±0.04 | 1.00±0.04 |
| + 490 ng/kg/min AngII[2] | 167 | **1.38** | **0.93**±0.04 | **2.62**±0.19[1] | 0.83±0.01[1] | **0.50**±0.05[1] | **0.92**±0.04 | **0.27**±0.05[1] |
| @ higher *P* | 167 | **1.38** | **1.07**±0.01 | **0.94**±0.01 | 1.00±0.01 | **1.57**±0.03 | **1.83**±0.06 | **1.32**±0.05 |
| C57;129SvEv$_{Fbln5}$[3] | 120 | 1.00 | 1.00±0.02 | 1.00±0.01[2] | 1.00±0.01[2] | 1.00±0.03[2] | 1.00±0.05 | 1.00±0.05[2] |
| + 490 ng/kg/min AngII[4] | 160 | **1.33** | **1.21**±0.01[1][2][3] | **1.06**±0.06[2] | 1.05±0.01[2] | **1.54**±0.08[1][2][3] | **1.52**±0.08[1][2][3] | **1.50**±0.08[1][2][3] |
| @ higher *P* | 160 | **1.33** | **1.06**±0.03 | **0.95**±0.01 | 1.00±0.01 | **1.50**±0.06 | **1.74**±0.09 | **1.31**±0.08 |
| C57;$Apoe^{-/-}$[5] | 112 | 1.00 | 1.00±0.02[4] | 1.00±0.03[2] | 1.00±0.01[2] | 1.00±0.02[2][4] | 1.00±0.04[4] | 1.00±0.01[2][4] |
| + 1000 ng/kg/min AngII[6] | 181 | **1.62** | **1.03**±0.05[4] | **1.61**±0.32[2] | 0.94±0.02[2][4] | **1.17**±0.18[2] | **1.48**±0.14[1][2][3][5] | **0.93**±0.18[2][4] |
| @ higher *P* | 181 | **1.62** | **1.10**±0.02 | **0.92**±0.02 | 1.00±0.01 | **1.93**±0.03 | **2.44**±0.10 | **1.52**±0.03 |

Mean±standard error values of systolic blood pressure plus the associated inner diameter and wall thickness, in vivo value of axial wall stretch, circumferential wall stress and material stiffness, and elastically stored energy, all as a ratio with respect to the respective normotensive groups; absolute systolic pressure [mmHg] was added for reference (first column). Data are from three previous studies: Bersi et al. 2016[5] (C57BL/6J, blood pressure measured using telemetry), Spronck et al. 2020[7] (C57;129SvEv$_{Fbln5}$), and Bersi et al. 2017[6] (C57;$Apoe^{-/-}$; blood pressure measured using tail-cuff method). The C57;129SvEv$_{Fbln5}$ mice were generated by breeding $Fbln5^{+/-}$ mice to obtain $Fbln5^{+/+}$ (i.e., wild-type) mice. $n$=5 for all groups. Superscripted numbers indicate statistically significant differences (one-way analysis of variance with post-hoc Bonferroni tests). AngII, angiotensin II. For discussion and absolute metric values, see Figure S7 and its legend.
